# Integrated phenomic and transcriptomic analyses unveil superior drought plasticity of North African durum wheat landraces

**DOI:** 10.64898/2026.04.03.716342

**Authors:** Rania Djemal, Rahma Trabelsi, Imen Ghazala, Chantal Ebel, Maxim Messerer, Rihab Boukouba, Maroua Gdoura-Ben Amor, Safa Charfeddine, Amine Elleuch, Radhouane Gdoura, Klaus F. X. Mayer, Jana Barbro Winkler, Jörg-Peter Schnitzler, Moez Hanin

## Abstract

Drought is a major constraint on the productivity of durum wheat across Mediterranean and North African regions. To elucidate the mechanisms underlying drought resilience, we employed a combination of scenario-controlled phenomics and flag leaf transcriptomics across ten durum wheat genotypes. These included the Tunisian landraces Chili and Mahmoudi, seven breeding lines, and the reference cultivar Svevo. The plants were grown to maturity under well-watered or long-term drought conditions in pots and rhizotrons, enabling a comprehensive assessment of growth, yield components, root architecture, physiological traits, and reaction norm plasticity. Drought markedly reduced performance, yet Chili and Mahmoudi consistently maintained superior biomass, grain number and intrinsic water use efficiency (iWUE). This was supported by balanced C/N allocation, strong osmotic adjustment, and the ability to sustain robust root systems under stress, albeit through partly divergent physiological strategies. Transcriptomic profiling revealed highly genotype specific responses, with drought tolerance unrelated to the number of differentially expressed genes. Instead, the landraces displayed distinct regulatory programs involving mainly photosynthesis protection, ABA-related transporters, osmotic adjustment pathways, and stress-responsive transcription factors. These mechanistic insights identify actionable physiological and molecular determinants of drought plasticity and provide high value targets for accelerating the breeding of climate resilient durum wheat.

**Highlights:** Integrated phenomics and transcriptomics revealed landrace-specific physiological and molecular mechanisms enabling superior drought resilience and identifying actionable targets for durum wheat improvement.

## Introduction

The Green Revolution of the mid-20th century markedly increased global food production through the deployment of semi-dwarf high-yielding crop varieties, together with synthetic fertilizers, pesticides, and improved irrigation practices (Xiao *et al.,* 2025). However, this progress came at the cost of reduced agrobiodiversity: diverse, locally adapted landraces were largely replaced by a limited number of modern cultivars, leading to a strong genetic erosion (Pingali, 2012; Lopes *et al*., 2015). In the context of climate change, the reduced diversity of high-yielding varieties has amplified their vulnerability, resulting in greater sensitivity to abiotic stresses (i.e. drought and heat) and to pests and diseases (Adhikari *et al*., 2022; Babalola *et al*., 2025). This situation urges scientists and breeders to accelerate breeding programs for the selection of more climate-resilient crops, notably by reintroducing local landraces as sources of adaptive alleles and by integrating yield potential with improved stress tolerance.

Durum wheat (*Triticum turgidum* ssp*. durum*) is a crucial cereal crop in Mediterranean Basin and North Africa and a key contributor to global food security (Enghiad *et al*., 2017). Yet this Mediterranean-North African belt is simultaneously recognized as a climate change hotspot for wheat production (Zampieri *et al*., 2017, 2020). In recent years, drought and heat have emerged as the primary abiotic stresses limiting global durum wheat production, particularly in arid and semi-arid regions such as North Africa (Aberkane *et al*., 2021; Melki *et al*., 2024). Water availability is essential across growth stages from early development to grain filling (Abid *et al*., 2018), and drought stress occurring during the vegetative phase can significantly reduce wheat grain yield (Sehgal *et al*., 2018; Cohen *et al*., 2021). Therefore, it is more than ever necessary to invest in breeding for the selection of new drought-tolerant durum wheat varieties.

Leveraging existing landraces offers a particularly promising avenue, because these populations have been shaped by long-term farmer selection in local environments and often retain allelic variations for stress adaptation that are rare or absent in elite pools (Lopes *et al.,* 2015; Broccanello *et al.,* 2023). This is especially relevant for durum wheat because drought tolerance is polygenic, scenario-dependent and often involves trade-offs between growth maintenance, water use regulation, carbon allocation and reproductive stability. As emphasised by the ‘drought-scenario’ framework, meaningful genetic gain requires access to broad diversity and phenotyping strategies that capture relevant drought profiles (timing, intensity and dynamics), rather than relying on single end-point measurements (Tardieu, 2012).

In Tunisia, durum wheat landraces were previously reported to be genetically diversified (Medini *et al*., 2005; Robbana *et al*., 2019; Slim *et al*., 2019). The two landraces ‘Chili’ and ‘Mahmoudi’ are among those introduced in Tunisia during early breeding programs by French colonists in the early 20th century. Both landraces are particularly interesting, as they have evolved naturally and have been selected by farmers over many generations for their disease resistance and adaptation to Mediterranean climatic conditions (Boukid *et al*., 2018). Comparative analyses have identified loci that differ between Tunisian landraces and modern cultivars, precisely the kind of variation required to expand elite breeding pools and identify markers associated with resilience (Robbana *et al*., 2019, 2021; Miazzi *et al*., 2022). More broadly, Mediterranean-wide analyses further support the existence of regionally structured alleles in durum landraces affecting yield components and phenology, reinforcing the value of landraces as sources of actionable marker variation (Soriano *et al*., 2018). Understanding these adaptive responses requires advanced methodologies, as drought tolerance involves various physiological and phenological processes influenced by multiple genetic factors and individual plant sensitivity to microenvironmental conditions (Roitsch *et al*., 2019; Zhao *et al*., 2019). Crop phenomics and high-throughput phenotyping (HTP) enable the repeated, non-destructive measurement of plant growth, architecture, and physiology on a large-scale using imaging and sensor-based pipelines (Cobb *et al*., 2013; Roitsch *et al*., 2019; Zhao *et al*., 2019; Yang *et al*., 2020). Such systems integrate data acquisition hardware, control terminals, and analytical software to enable non-destructive, high-throughput evaluation of observable and non-observable plant traits (Xiao *et al*., 2022; Alptekin, 2024). These methods are relevant alternatives to the labor-intensive nature of traditional techniques, enabling continuous monitoring of plant reactions under stress-controlled conditions and improved understanding of plant-environment interactions (Zhao *et al*., 2019). In wheat, the recent focus has shifted towards ‘functional phenotyping’, where physiological traits (and their dynamics) are quantified under defined stress scenarios to strengthen causal inference and deliver actionable traits for breeding (Correia *et al*., 2022). Controlled-environment platforms are particularly powerful for drought studies because they enable reproducible scenarios and high temporal resolution. For instance, imaging combined with physiological readouts has already been shown to effectively discriminate contrasting early-stage water-use strategies in durum wheat genetic resources (Nakhforoosh *et al*., 2016). However, extracting breeding-relevant value from phenomics depends critically on (i) appropriate scenario design and (ii) robust integration methods that can interpret genotype × environment interactions and trait covariance structures across time and organs (Tardieu, 2012; Coppens *et al*., 2017; Yang *et al*., 2020). More importantly, integrating HTP with molecular profiling combines quantitative, time-resolved measurements of plant performance and plasticity under stress with transcriptomic insights. Such integration enables the identification of regulatory programs and pathway reconfigurations underlying phenotypic alterations, thereby strengthening the genotype-to-phenotype chain. This helps to partition drought responses into physiological modules (e.g. growth maintenance vs. water-use regulation); identify candidate regulators and pathways associated with superior performance; and prioritise robust markers for selection using data-driven integration approaches (Coppens *et al*., 2017; Zhao *et al*., 2019; Yang *et al*., 2020). In durum wheat, these integrative strategies are becoming more feasible as high-quality genomic resources, notably the ‘Svevo’ cultivar reference genome, provide a foundation for transcriptome interpretation and marker development (Maccaferri *et al*., 2019). In parallel, recent genetic studies have begun mapping genomic regions underlying drought-relevant productivity traits, helping in translating mechanistic candidates into breeding targets (Zaïm *et al*., 2024).

Despite this promise, studies that combine high-resolution phenomics drought responses with genotype-specific transcriptional programs in North African durum landraces are limited, particularly under controlled drought scenarios that enable mechanistic analysis and robust cross-genotype comparison. Here, we use an integrated phenomics–transcriptomics strategy to characterise the drought responses of ten durum wheat genotypes including the Tunisian landraces ‘Chili’ and ‘Mahmoudi’, alongside improved cultivars and the reference genotype ‘Svevo’. Our approach combines controlled-environment phenomics under a defined water-deficit scenario to quantify agronomically relevant growth and yield traits and physiological indicators, with RNA-seq profiling of flag leaves to resolve genotype-specific transcriptional reprogramming. We aim to identify phenotypic indicators that best discriminate against drought-tolerant performance by explicitly quantifying phenotypic plasticity (reaction norms) across key traits and linking these responses to transcriptional programs. Specifically, we address four key questions in this study. First, we examine how drought-induced phenotypic plasticity differs among Tunisian landraces, improved lines, and the reference cultivar ‘Svevo’ across growth, yield, and major physiological traits. Second, we identify which phenomic and physiological traits most effectively discriminate drought-tolerant performance under a defined long-duration water-deficit scenario. Third, we test whether the landraces ‘Chili’ and ‘Mahmoudi’ achieve superior drought performance through shared or contrasting physiological strategies, particularly in relation to water-use regulation, osmotic adjustment, and resource allocation. Finally, we investigate which flag-leaf transcriptional programs and regulatory pathways drive drought plasticity and identify genes that can be translated into breeding markers for Mediterranean and North African environments.

## Materials and Methods

### Plant Material

A total of ten durum wheat (*Triticum turgidum* ssp. *durum*) genotypes, provided by the National Institute of Field Crops, Tunisia, were used, including the reference cultivar ‘Svevo’ and nine Tunisian varieties. The latter included two traditional landraces (‘Mahmoudi’ and ‘Chili’) and seven improved breeding lines (‘Khiar’, ‘Salim’, ‘Maali’, ‘Razzek’, ‘Karim’, ‘Dhahbi’ and ‘INRAT100’). The seeds were surface-sterilised in a solution of 5% sodium hypochlorite for 10 minutes, rinsed three times with sterile distilled water and then germinated for 3 and 5 days on moist filter paper in Petri dishes before being transplanted into pots and a rhizotron system, respectively. Wheat growth was evaluated according to phenological growth stages defined by (Lancashire *et al.,* 1991).

### Experimental design and growth conditions

The plants were cultivated in the Fitness-SCREEN high-throughput phenotyping facility (Experimental Simulation Unit, Helmholtz Munich), which is a climate-controlled greenhouse with automated irrigation and environmental monitoring (Bose *et al.,* 2024; Newrzella *et al.,* 2025). Two cultivation systems were run in parallel (Supplementary Fig. S1). For shoot-focused phenotyping, single plants were grown in 8.5 L polypropylene pots. For combined above- and below-ground phenotyping, plants were grown in 400 × 800 × 40 mm rhizotrons that enabled non-invasive root imaging through a transparent observation window. A peat-based substrate (Einheitserde Classic type CL-T, Patzer Erden GmbH, Germany) mixed with black basalt sand (0.6–1.2 mm grain size) at a ratio of 1:1.2 (v/v) was used for both systems. In the rhizotrons, three plants were established per unit and spaced 100 mm apart to approximate a field-like density. Genotype identities were randomised across carriers/positions to minimise spatial biases.

Drought treatment was initiated at the BBCH 30 stage and continued until physiological maturity (BBCH 95). While control plants were maintained at 70% net pot capacity, drought-stressed plants were brought to and maintained at 30%. To achieve the target level of drought stress, irrigation was withheld until the substrate water content declined; accordingly, thereafter, irrigation was adjusted to maintain this level. Substrate water content (net pot capacity (%)) was monitored gravimetrically throughout the experiment, in line with established methods for estimating pot capacity and related reference points in soils/substrates (Grewal *et al*., 1990). There were n = 9 plants per genotype per treatment in the pot system and n = 5 rhizotrons per genotype per treatment in the rhizotron system (three plants per rhizotron).

### Non-invasive phenotyping

Plants were regularly monitored throughout their development under both irrigation regimes. At each weighing, a 2D image of the plants was captured using an RGB camera. These images were then used to derive plant-specific traits (such as height and number of leaves). In the rhizotron system, non-invasive below-ground phenotyping was performed by imaging the transparent side of each rhizotron with a fixed camera setup (Allied Vision Prosilica GT 6600 with a Milvus 2/50 M lens). Images were acquired as grayscale files spanning the full 685 × 322 mm observation window (approximately 8.8 pixels per mm) at a resolution of 6012 × 2821 pixels. Root image analysis followed an AI-assisted segmentation and trait extraction workflow. The raw images were converted from .bmp to .tif format and then segmented using RootPainter (v0.2.21). This software employs convolutional neural networks and an iterative corrective-annotation training strategy to refine model predictions (Smith *et al*., 2022). The segmented masks were then imported into RhizoVision Explorer (v2.0.3) for the quantitative extraction of root traits (Seethepalli *et al*., 2021). In RhizoVision Explorer, “broken roots” mode was enabled to improve the reconstruction of discontinuous segments and uniform filters were applied across batches (threshold = 200, removal of non-root objects >1 mm², root pruning threshold = 1). The primary dynamic root traits reported here were total visible root length, root surface area and number of root tips, extracted for each image and time point.

### Destructive harvest traits, yield components, and drought indices

At the BBCH 95 stage, the plants were harvested for destructive measurements. The following above-ground biomass and yield components were quantified: total plant biomass; ear number per plant; ear weight; grain number per plant; grain yield per plant; and thousand-kernel weight (TKW).

To evaluate the drought tolerance, six indices were calculated using grain yield per plant (rhizotron system) under control (Yp) and drought (Ys) conditions: mean productivity (MP) (Rosielle and Hamblin, 1981); geometric mean productivity (GMP) (Fernandez, 1992); tolerance index (TOL) (Rosielle and Hamblin, 1981); yield stability index (YSI) (Bouslama and Schapaugh, 1984); stress susceptibility index (SSI) (Moosavi *et al*., 2007) and stress tolerance index (STI) (Fernandez, 1992) The indices were calculated as follows:

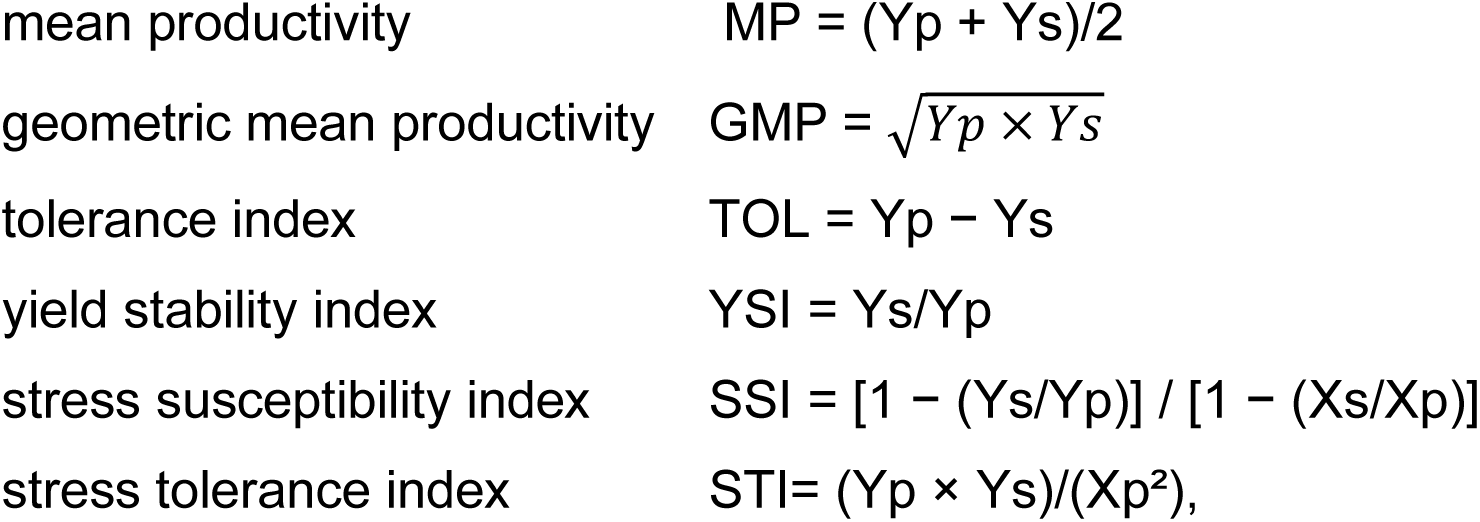

where Yp and Ys are the yield performance of varieties, while Xp and Xs are the mean yield of all varieties under control and stress, respectively (Lamba *et al*., 2023).

To measure phenotypic plasticity, we quantified the magnitude of the plastic response using the relative distance plasticity index (RDPI), ranging from 0 (no plasticity) to 1 (maximal plasticity), by RDPI = (|Xc —Xs|)/(|Xc + Xs|), where Xc and Xs represent the different trait values of biological replicates under control and stressed conditions (Xiao *et al*., 2025).

### Stable isotope and elemental analyses and intrinsic water-use efficiency

The elemental concentrations of carbon (C) and nitrogen (N) and the carbon isotope composition (δ¹³C) were determined in leaf material that had been separated into soluble and solid fractions. The material was analysed using an elemental analyser (Euro EA, Eurovector, Milan, Italy) coupled to an isotope ratio mass spectrometer (IRMS; Delta V Advantage, Thermo Fisher, Dreieich, Germany). This was done in accordance with the workflow and reporting conventions described by Nosenko *et al.,* (2025). The carbon isotope composition was expressed relative to the VPDB standard as follows:

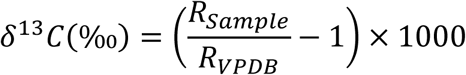

where R_sample_ is the ¹³C/¹²C ratio of the sample and R_VPDB_ = 0.0111802 (Werner and Brand, 2001).

Intrinsic water-use efficiency (iWUE) was derived from δ¹³C values using the R package isocalcR (Mathias and Hudiburg, 2022), in line with the approach outlined by (Nosenko *et al*., 2025). The mean air temperature over the leaf development period (23/05/2024–30/07/2024) was obtained from the daily records of German Weather Service (DWD) station 1975. The atmospheric CO₂ isotope composition inputs were taken from the published compilations described in (Nosenko *et al*., 2025).

### Leaf Osmotic Potential

The leaf osmotic potential (π) was quantified using the extraction and osmometry approach described by (Nosenko *et al*., 2025). In brief, 10 mg of dried leaf tissue powder was suspended in 750 µL of 10 mM potassium phosphate buffer solution (KPi), pH 7.0. Osmolarity was measured from a 50 µL sample using a Röbling Type 13 freezing-point osmometer (Röbling Messtechnik GmbH, Berlin). Buffer osmolarity was subtracted from sample readings. The osmotic potential (π) was then calculated using the van’t Hoff equation, as described by (Nosenko *et al*., 2025). This equation incorporates osmolarity, the universal gas constant, absolute temperature, and the ratios of dry mass to extraction volume, as well as the ratio of fresh-to-dry mass.

### RNA sequencing and differential expression analysis

Flag leaves were sampled under control and drought conditions at the reproductive stage (BBCH 75; Supplementary Fig. S1), immediately frozen in liquid nitrogen and stored at −80°C until processing. Total RNA was extracted using a phenol–chloroform method based on a guanidinium isothiocyanate extraction buffer (TriReagent). Approximately 50–100 mg of frozen flag leaf tissue was ground into a fine powder in liquid nitrogen. The powder was homogenized in 1 mL of RNA extraction buffer and incubated on ice (≈8°C) with vigorous mixing. Subsequently, 400–500 µL of ice-cold chloroform were added, and samples were mixed by inversion before incubation at room temperature. Following centrifugation (13,000 rpm, 4°C, 10–12 min), the aqueous phase was carefully transferred to a new RNase-free tube. To remove genomic DNA contamination, samples were treated with RNase-free DNase I at 37°C for 30 min, followed by a second chloroform extraction and centrifugation. RNA was precipitated with ice-cold isopropanol, incubated on ice, and centrifuged to obtain the RNA pellet. The pellet was washed twice with 70% ethanol, air-dried, and resuspended in RNase-free water. Then, RNA quality was assessed using a NanoDrop spectrophotometer (Thermo Fisher Scientific). RNA samples were immediately stored in −80°C and subsequently sent for further processing, library preparation and sequencing. Paired-end (150 bp × 2) reads of control and treated samples were generated using Illumina NovaSeq X Plus. Library preparation was done by Poly-A enrichment. The quality of the RNA-seq reads was assessed using FastQC (Andrews, 2010). Adapter trimming was performed using Cutadapt (Martin, 2011). Transcript abundance was quantified by pseudo-alignment to the durum wheat reference transcriptome Svevo v1.0 (Maccaferri *et al*., 2019) using Kallisto (Bray *et al*., 2016). Differential gene expression analysis was conducted in DESeq2 (Love *et al*., 2014). Genes were considered to be differentially expressed at a false discovery rate (FDR) of less than 0.05, with an additional fold-change threshold applied where indicated in the Results section. Mercator4 (Bolger *et al*., 2021) was used for functional annotation of the Svevo v1.0 transcriptome. The resulting BINs were analyzed separately, genes which were not significantly differentially expressed in any sample were discarded, and the mean logFC of each BIN per sample was calculated. The results were plotted as heatmaps using the R-package pheatmap (Kolde, 2025). UpSet plots were generated using the R-package UpSetR (Conway *et al*., 2017).

### Statistical analyses

The graphs were produced using GraphPad Prism 10.1.0 software. Statistical analyses were carried out using SAS JMP Pro 17 software. Treatment effects were evaluated using analysis of variance (ANOVA), with mean separation performed via Tukey’s test at P < 0.05. Pearson correlation coefficients were calculated to evaluate the relationships between the measured traits and the drought indices. Significance was assessed at P < 0.05, P < 0.01 and P < 0.001. Multivariate analyses (PCA and PLS-DA) were performed in MetaboAnalyst 6.0 (Pang *et al*., 2024) after the data were log-transformed.

## Results

### Yield- and growth-related traits discriminate against performance under drought

We used a high throughput phenotyping approach to perform a comparative assessment of above-ground traits of 10 durum wheat genotypes growing under control (70% of net pot capacity (PC)) or drought conditions fixed as 30% net PC from BBCH 30 to the reproductive stage (BBCH 95, see Supplementary Fig. S1). Out of these ten varieties, the well characterized and fully sequenced genotype Svevo was used as a reference (Maccaferri *et al*., 2019). The remaining nine durum wheat genotypes are locally grown in Tunisia and made of two landraces (Chili and Mahmoudi) and seven improved lines (Khiar, Salim, Maali, Razzek, Karim, Dhahbi and INRAT100). Under controlled greenhouse conditions, the experimental setup was based on two parallel cultivation systems, in which the plants were either placed individually in pots or in sets of three plants in a rhizotron (see Materials and Methods).

As expected, all genotypes exhibited significant reductions of the different parameters measured under drought stress, compared with control conditions, in both pot (Supplementary Fig. S2, S3) and rhizotron systems (Fig. 1). Most of the growth and yield parameters observed on plants grown in pots correlated relatively well with those grown in rhizotrons, as confirmed by the correlation matrix based on the data collected for yield and growth under control and drought stress conditions (Supplementary Fig. S4). Such results emphasize the impact of drought on the durum wheat genotypes in different growth environments and confirm the robustness of the data. Therefore, we will hereafter mostly discuss the data retrieved from plants grown in rhizotron as it reflects better the field growth conditions (Cope *et al*., 2024) since plants are submitted to additional competition for resources which is due to space occupied by the neighbouring plants in the same rhizotron.

**Figure 1:**
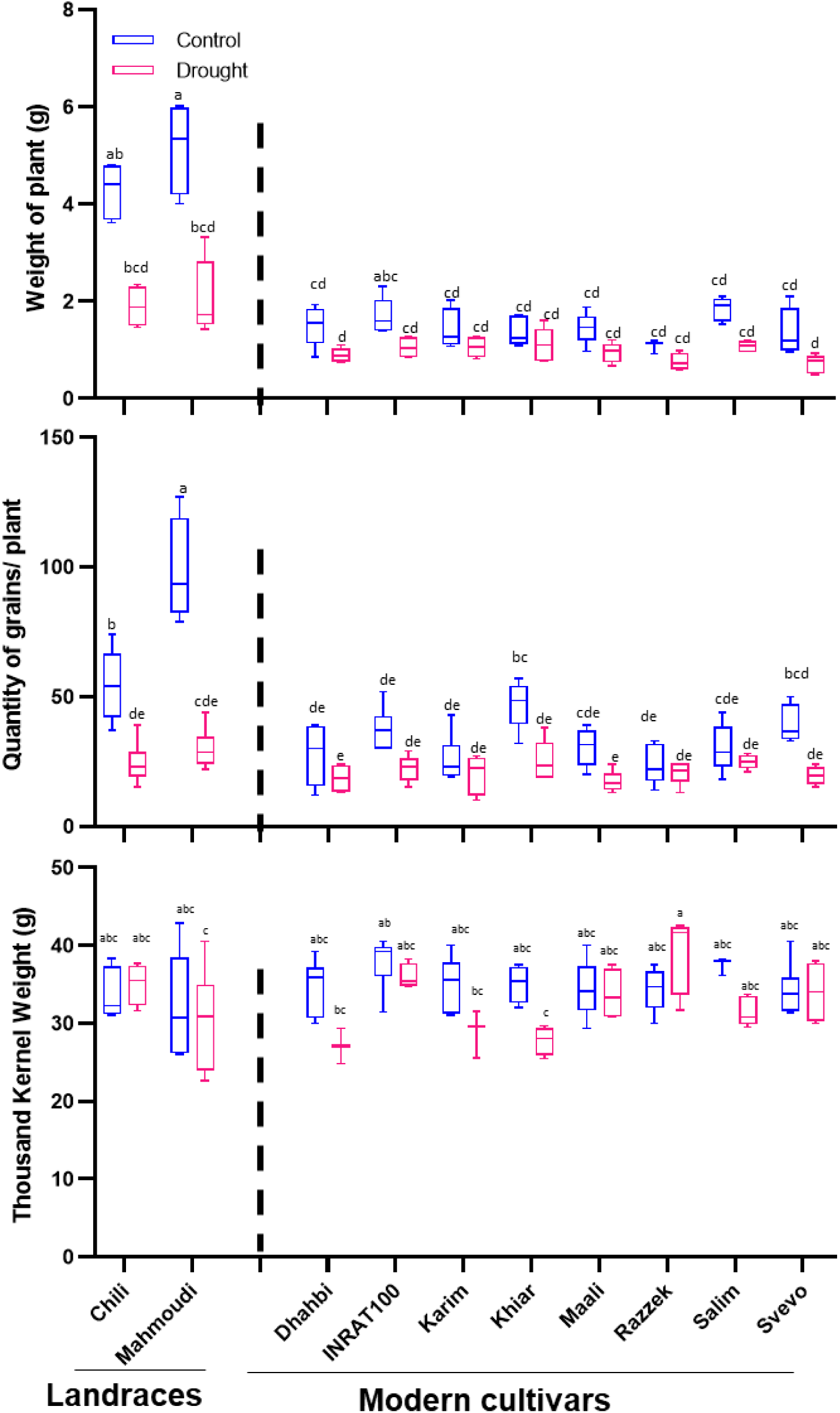
Yield perfomance of 10 durum wheat genoytpes under control and drought stress cultivated in rhizotrons at maturity (BBCH 95). A) Weight of the plants in g. B) Quantity of grains/plant C) Thousand Kernel Weight (g). Data represent the means and standard deviations (error bars) of six biological replicates of plants grown in rhizotrons or pots. Different letters indicate significant differences (P < 0.05) among samples, as determined by one-way ANOVA followed by Tukey’s HSD test.

Plant weight is an important agronomic trait often associated with enhanced straw and grain yields (Chamekh *et al*., 2015). At BBCH 95 under control conditions, the two landraces Chili and Mahmoudi had the greatest plant biomass (approx. 4 g) while for the other lines, it ranged from 2 to 1.5 g. The drought stress negatively impacted the growth to different extents depending on the genotypes. For instance, genotypes like Khiar, Karim, Razzek and Dhahbi were weakly affected by drought stress, whereas Maali, INRAT100 and Svevo showed a moderate biomass reduction. Interestingly, although a significant decrease in biomass of both landraces (50% and 60% reduction for Chili and Mahmoudi, respectively) was observed, they seem to withstand well the drought stress as they maintain higher biomasses compared to the other genotypes.

This trend was confirmed by measuring other yield traits such as the grain quantity per plant (Fig. 1B) and weights of ears and grains (Supplementary Fig. S2). The grain quantity per plant decreased for all genotypes under drought stress, but the landraces Chili and Mahmoudi exhibited the greatest yields with more than 25 grains per plant while this value was less than 20 for the other genotypes. Moreover, the measured biomass was positively correlated (p < 0.01) with the quantity of grains/plant (Supplementary Fig. S4). The calculation of thousand kernel weight (TKW) showed that Chili and Razzek genotypes exhibited higher TKW values under drought. In contrast, Khiar, Karim, and Dhahbi exhibited the most significant decrease in TKW under drought (Fig. 1C).

Similar scenario occurred when measuring root surface area and plant height (Supplementary Fig. S5), with the two landraces retaining the highest values compared to modern cultivars under control and drought conditions. The use of rhizotrons enabled the measurement of root parameters (total root length, root surface, and number of root tips) which revealed that drought has a genotype-dependent effect on root architecture with a clear a GxE interaction (Fig. 2, Supplementary Fig. S5). While Mahmoudi and Chili total root length was reduced at BBCH 60 under drought (−26% and −8% respectively), Dhahbi and Svevo maintained similar root length at this stage under drought. Svevo also exhibited the strongest increase in total root length (about 50%) between BBCH 60 and BBCH 75 under drought (Fig. 2). However, an increase in root branching and surface area was observed signaling compensatory fine-root proliferation under stress. This was especially true for Khiar, Maali and Svevo for which there were 145%, 114% and 75% more root tips at BBCH 75 compared to BBCH 60 under drought stress (Fig. 2, lower panels). Interestingly, Mahmoudi increased its root branching already at BBCH 60 under drought compared to BBCH 30 (+134% increase in root tips number under drought vs 54% increase compared to BBCH 30 under control conditions) and lost 10% of its root tips at BBCH 75 under drought. Chili increased its root branching also at BBCH 75 compared to BBCH 60 by 92%. Therefore, the different genotypes seem to adopt diverse strategies for root development under drought to explore the soil for water.

**Figure 2:**
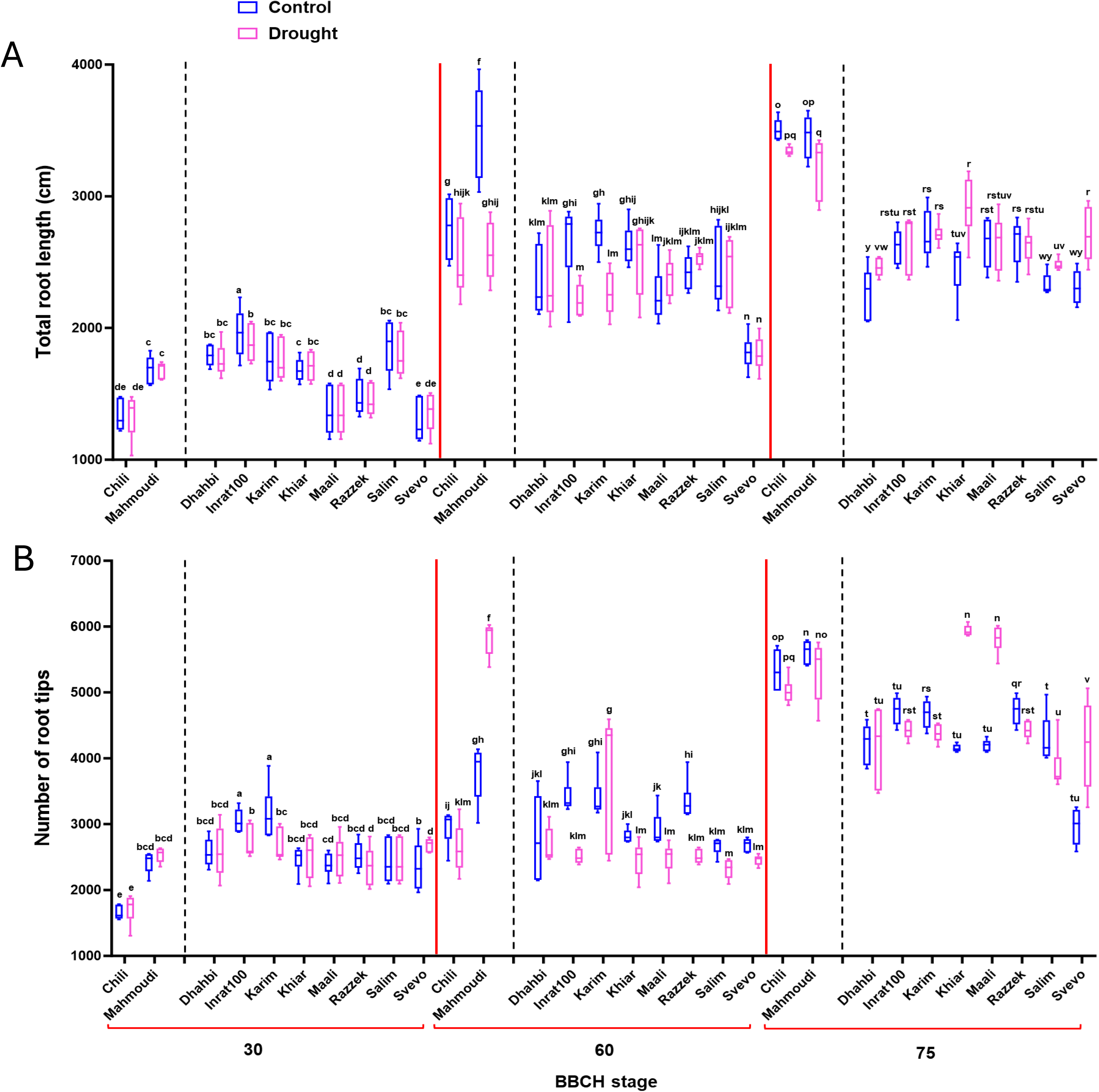
Comparative assessment of root traits: number of total root length (A), and number of root tips (B) at BBCH 30, BBCH 60, and BBCH 75 under control (blue boxes) and drought (pink boxes) conditions. Measurements were obtained using RhizoVision Explorer from segmented outputs generated by RootPainter based on images of roots from plants grown in the rhizotron. Box plots represent mean values ± SD (n = 6). GraphPad Prism 10.1.0 software was used for data visualization. Statistical analyses were carried out using SAS JMP Pro 17 software. Phenological stages: BBCH 30: Stem elongation; BBCH 60: onset of flowering; BBCH 75: grain development.

### Drought indices and multivariate ranking of genotypes

To have an overview of the behavior of the two landraces and the breeding lines towards drought, we evaluated six stress tolerance indices (Table 1, Fig. 3). The mean productivity (MP) provides information about how a genotype combines yield performance and stability as it highlights the one which is productive when water is available and able to maintain acceptable yield under drought. Whereas geometric mean productivity (GMP) is considered as one of the most robust indicators of drought tolerance as it takes into consideration productivity, stability and drought tolerance. According to the mean productivity (MP) and the geometric mean productivity (GMP) indices, the two landraces exhibit the highest values, followed by INRAT100. This trend seems to correlate with the stress tolerance index (STI), where the values registered exceed 1 (i.e. 1.51 for Mahmoudi and 1.28 for Chili), synonyms of drought-tolerant and high yielding varieties. However, these parameters are not sufficient to qualify these three varieties as the most drought tolerant and must be interpreted in association with other indices. For instance, the Tolerance Index (TOL), commonly used in plant breeding, measures a genotype’s ability to maintain yield under stress conditions. Based on our data, Razzek exhibited the lowest TOL values, but the two landraces ranked the last. Considering the Yield Stability Index (YSI), the breeding lines Razzek and Maali were the most tolerant genotypes, since they maintained yield closest to their non-stressed potential. The Stress Susceptibility Indices (SSI) varied from 0.21 for Razzek (least susceptible) to 1.29 for Svevo and Chili. An objective evaluation of drought tolerance should consider both productivity (MP, GMP, STI) and stability (TOL, YSI, SSI), leading to the following ranking: Mahmoudi > Chili > INRAT100 > Salim > Razzek > Maali > Svevo > Khiar > Karim > Dhahbi. This ranking is well illustrated in the PCA biplot based on the six indices, where the two landraces cluster together followed by INRAT100, while Razzek, Maali, Karim and Dhahbi occupy the most distant and opposing positions (Fig.3).

**Figure 3:**
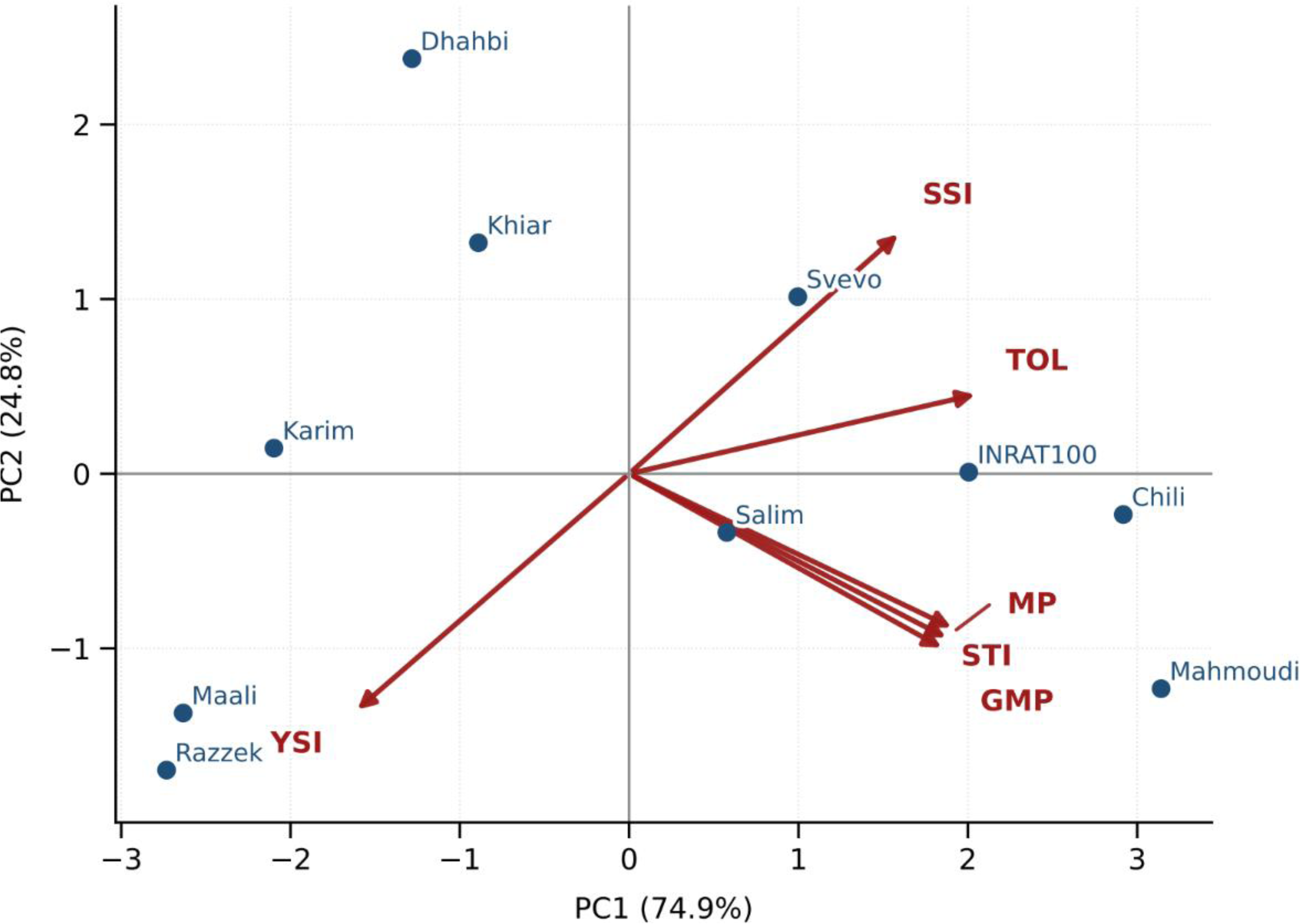
PCA biplot of drought tolerance and productivity indices. The biplot PCA scores for ten wheat varieties (blue points with labels) and variable loadings (arrows) computed from six indices: MP (mean productivity), GMP (geometric mean productivity), STI (stress tolerance index), SSI (stress susceptibility index), YSI (yield stability index), TOL (tolerance index). Vectors represent correlation loadings of each index on the same components. PC1 and PC2 explain 74.9% and 24.8% of the total variance, respectively.

**Table 1:**
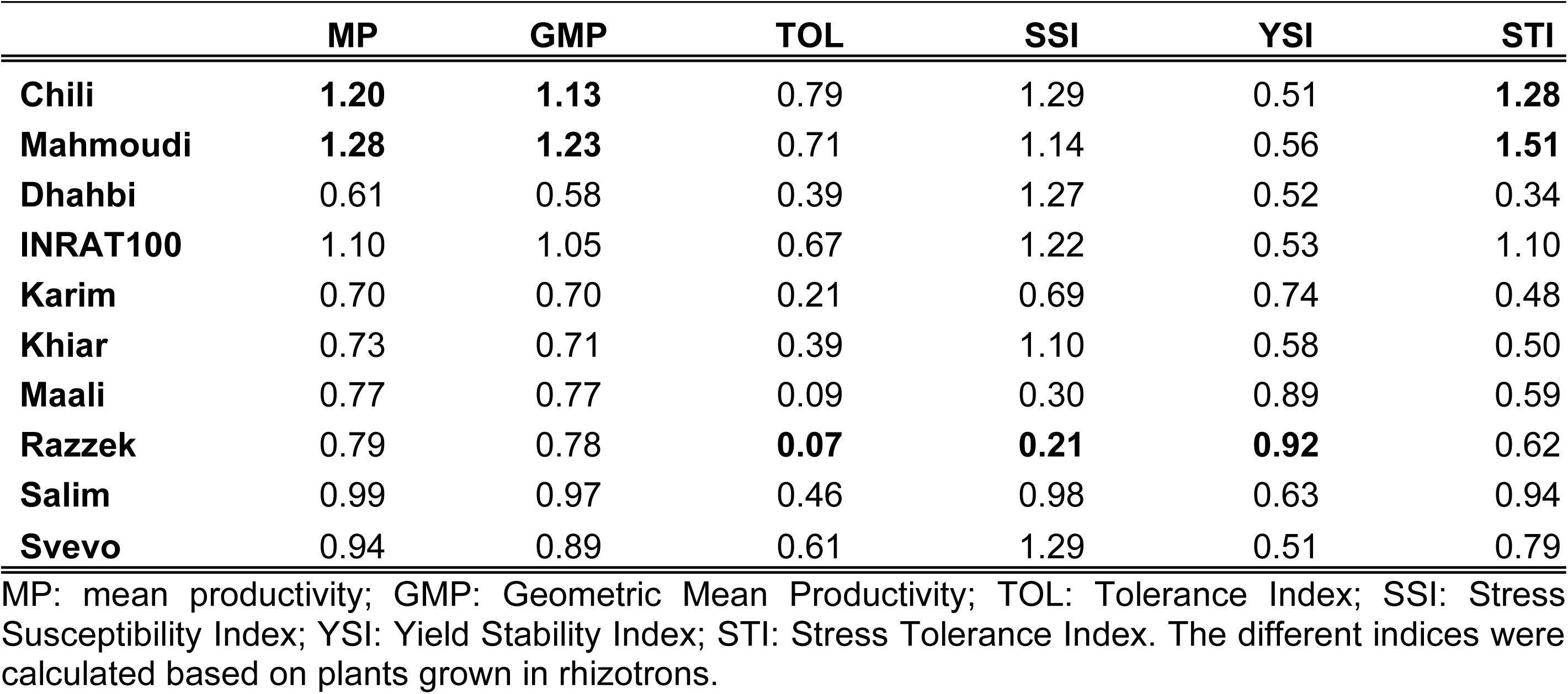
Influence of drought conditions on yield and different indices.

The observed genotype by environment interaction underscores the critical importance of multi-environment screening to accurately assess drought tolerance/performance and select the most promising genetic material for durum wheat breeding programs aimed at enhancing climate resilience. Based on the different growth and yield parameters, it seems that both landraces rank among the drought-tolerant and high-performing durum wheat varieties.

### Phenotypic plasticity quantified by reaction norms

The comprehensive phenotypical analysis of this core durum wheat collection revealed genetic variation of their drought response with the two landraces behaving as best performing under water stress. This finding prompted us to assess the phenotypic plasticity in drought response, considering several traits including plant weight, weight of ears and grains, quantity of grains/plant, TKW, total root length, root surface, number of root tips. This phenotypic plasticity was measured using the relative distance plasticity index (RDPI) (Xiao *et al.,* 2025). As shown in Table 2, we observed a high variability in the RDPI values among the wheat genotypes fluctuating between 0.01-0.05 (almost no plasticity) and up to 0.43 (the maximal observed plasticity). Based on these traits, Razzek, followed by Maali seem to be the least flexible, while the two landraces exhibit the highest plasticity particularly for yield traits. Interestingly, the reference cultivar Svevo displayed phenotypic plasticity for both above and below-ground traits.

**Table 2:** Relative Distance Plasticity Index values for above and below-ground phenotypic traits.

| Traits | Chili | Mahmoudi | Dhahbi | INRAT100 | Karim | Khlar | Maali | Razzek | Salim | Svevo |
| --- | --- | --- | --- | --- | --- | --- | --- | --- | --- | --- |
| **Weight of the plant** | 0.37 | 0.25 | 0.14 | 0.22 | 0.10 | 0.09 | 0.12 | 0.05 | 0.17 | 0.23 |
| **Weight of grains** | 0.33 | 0.28 | 0.29 | 0.30 | 0.15 | 0.19 | 0.06 | 0.04 | 0.23 | 0.33 |
| **Weight of Ears** | 0.42 | 0.28 | 0.11 | 0.26 | 0.13 | 0.12 | 0.10 | 0.07 | 0.21 | 0.32 |
| **Qty of grains/plant** | 0.43 | 0.37 | 0.19 | 0.27 | 0.07 | 0.05 | 0.14 | 0.05 | 0.10 | 0.40 |
| **TKW** | 0.13 | 0.06 | 0.07 | 0.04 | 0.07 | 0.16 | 0.06 | 0.01 | 0.14 | 0.05 |
| **Root Tips Number** | 0.041 | 0.036 | 0.078 | 0.032 | 0.033 | 0.177 | 0.160 | 0.032 | 0.069 | 0.168 |
| **Total Root Length** | 0.024 | 0.046 | 0.047 | 0.019 | 0.025 | 0.085 | 0.023 | 0.030 | 0.044 | 0.079 |
| **Root surface Area** | 0.018 | 0.073 | 0.083 | 0.039 | 0.053 | 0.099 | 0.094 | 0.029 | 0.056 | 0.120 |

| Traits | Chili | Mahmoudi | Dhahbi | INRAT100 | Karim | Khlar | Maali | Razzek | Salim | Svevo |
| --- | --- | --- | --- | --- | --- | --- | --- | --- | --- | --- |
| Weight of the plant | 0.37 | 0.25 | 0.14 | 0.22 | 0.10 | 0.09 | 0.12 | 0.05 | 0.17 | 0.23 |
| Weight of grains | 0.33 | 0.28 | 0.29 | 0.30 | 0.15 | 0.19 | 0.06 | 0.04 | 0.23 | 0.33 |
| Weight of Ears | 0.42 | 0.28 | 0.11 | 0.26 | 0.13 | 0.12 | 0.10 | 0.07 | 0.21 | 0.32 |
| Qty of grains/plant | 0.43 | 0.37 | 0.19 | 0.27 | 0.07 | 0.05 | 0.14 | 0.05 | 0.10 | 0.40 |
| TKW | 0.13 | 0.06 | 0.07 | 0.04 | 0.07 | 0.16 | 0.06 | 0.01 | 0.14 | 0.05 |
| Root Tips Number | 0.041 | 0.036 | 0.078 | 0.032 | 0.033 | 0.177 | 0.160 | 0.032 | 0.069 | 0.168 |
| Total Root Length | 0.024 | 0.046 | 0.047 | 0.019 | 0.025 | 0.085 | 0.023 | 0.030 | 0.044 | 0.079 |
| Root surface Area | 0.018 | 0.073 | 0.083 | 0.039 | 0.053 | 0.099 | 0.094 | 0.029 | 0.056 | 0.120 |

For each trait, Relative Distance Plasticity Index (RDPI) was calculated using the following formula: $RDPI = (|X_c - X_s|) / (|X_c + X_s|)$ , where $X_c$ and $X_s$ represent the different trait values of biological replicates ( $n=6$ ) kept under control (c) and drought stress (s) conditions with 0 representing no plasticity and 1 maximal plasticity. Within each row (trait), greener colors correspond to higher RDPI values, whereas redder colors represent lower RDPI values.

### Physiological mechanisms underpinning drought performance and plasticity

#### Comparative Analysis of Leaf Nitrogen and Carbon partitioning in Tunisian Durum Wheat

Analysis of carbon (C) and nitrogen (N) partitioning (Fig. 4; Supplementary Fig. S7) provides insights into the physiological variations observed across the ten durum wheat genotypes in response to drought. Our results showed that compared to the other genotypes, Chili and Mahmoudi, have relatively higher levels of C and N in the solid fractions of their leaves suggesting their greater capacity to accumulate structural components (cell wall and photosynthetic proteins for insoluble C and N fractions, respectively). The integrative C/N ratio further delineated the drought responses among genotypes (Fig. 4). Sensitive varieties exhibited elevated C/N ratios under stress, whereas the ratio remained relatively balanced in the landraces. This contrast suggests that landraces are better able to maintain coordinated carbon assimilation and nitrogen metabolism under stress conditions.

**Figure 4:**
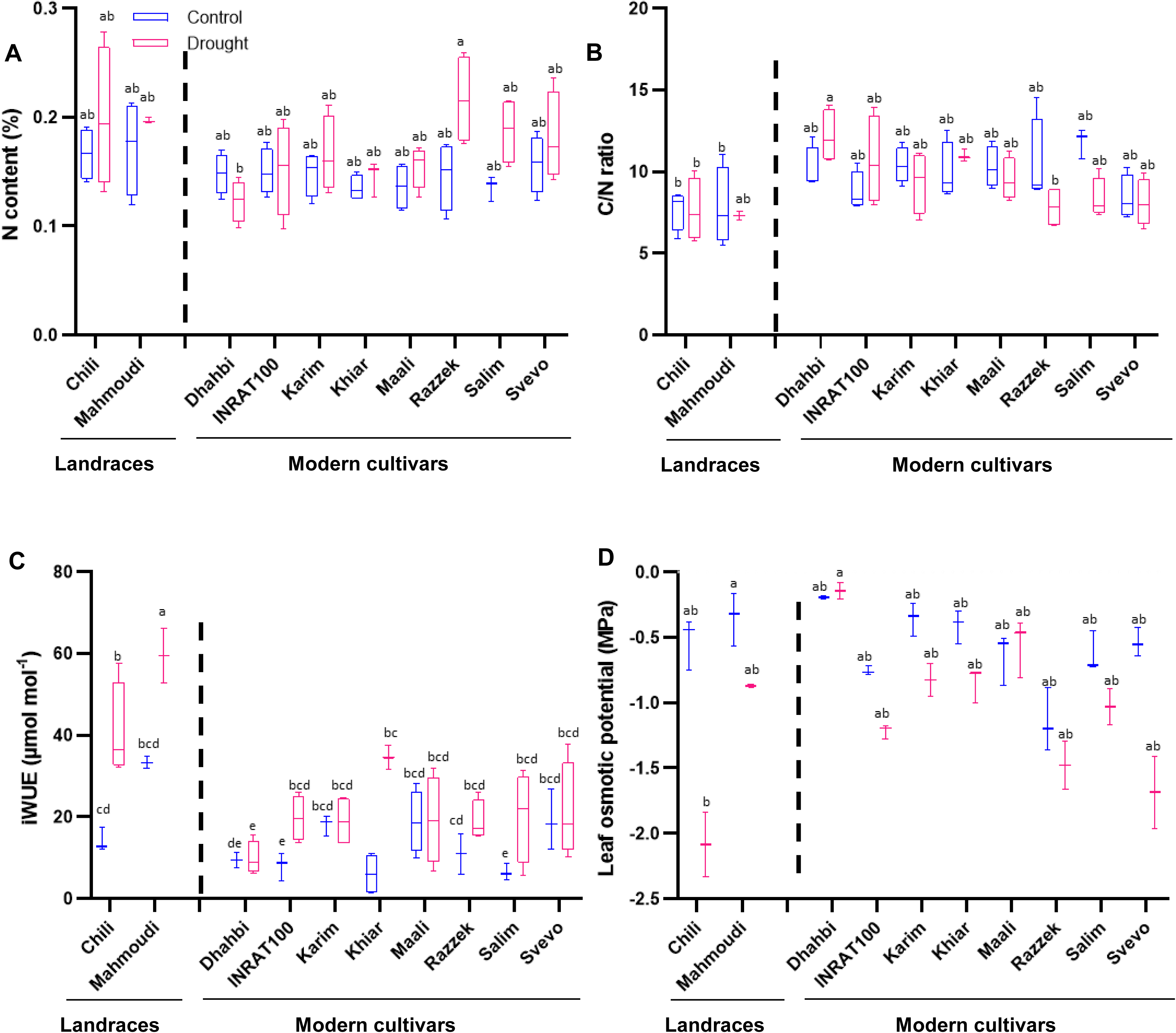
Physiological responses of ten durum wheat genoytpes to drought stress. Leaves (F-2, BBCH 75) of plants grown in rhizotrons were used to assess nitrogen content (A), and C/N ratio (B) from solid leaf samples across all varieties. (C) iWUE in solid fraction. (D) leaf osmotic potential. Data are mean values from four biological replicates. Different letters denote statistically differences between genotypes as determined by one-way ANOVA followed by Tukey’s HSD test (P < 0.05). Data are mean values from four biological replicates.

#### Leaf δ¹³C isotopic, intrinsic water-use efficiency and osmotic potential analyses

Further we measured the δ¹³C isotopic composition, which provides information on intrinsic water-use efficiency (iWUE) and carbon assimilation. We assessed the carbon partitioning patterns in solid and soluble tissue fractions under both controlled and water-stress conditions (Fig. 4 and Supplementary Fig. S8). Our results showed distinct response patterns between the two fractions. The solid fraction in the F-2 leaf, demonstrated genotype-specific stability in δ¹³C values under water deficit conditions indicating a variation in preserving carbon assimilation efficiency and structural integrity during drought stress. Compared to the breeding lines, Chili and Mahmoudi registered the highest δ¹³C values in the solid fraction, suggesting inherent preservation mechanisms characteristic of drought-tolerant genotypes. Consistently, the intrinsic water use efficiency (iWUE) was significantly higher in these two landraces (Fig. 4C), reflecting their greater capacity to fix carbon through photosynthesis while minimizing water loss, presumably via better control of stomatal closure (Lawson and Blatt, 2014). On the other hand, we noticed more pronounced drought-induced δ¹³C variations in the soluble fraction, reflecting active metabolic reorganization (Supplementary Fig. S8).

As a complementary drought-tolerance mechanism, osmotic adjustment (reflected by more negative leaf osmotic potential, Ψπ) often operating in parallel with iWUE, was also evaluated. Interestingly, Chili showed the most negative Ψπ values, suggesting that this variety may have a higher capacity to maintain cell turgor and water uptake under drought, most likely via stronger osmolytes (i.e. soluble sugars and proline) accumulation (Farquhar *et al*., 1989; Mininni *et al*., 2022) (Fig. 4D) and that the two landraces develop their stress resilience via different physiological routes.

A clustered heatmap combining the various phenotypical and physiological traits measured across the 10 durum wheat genotypes based on the data extracted from the two growth systems (pots and rhizotron; Fig. 5 and Supplementary Fig. S9), revealed that Mahmoudi and Chili consistently behave similarly for most of the parameters and cluster together both in control and drought stress conditions corroborating their similar phenotypic behavior, without ruling out the existence of distinct underlying drought-tolerance mechanisms.

**Figure 5:**
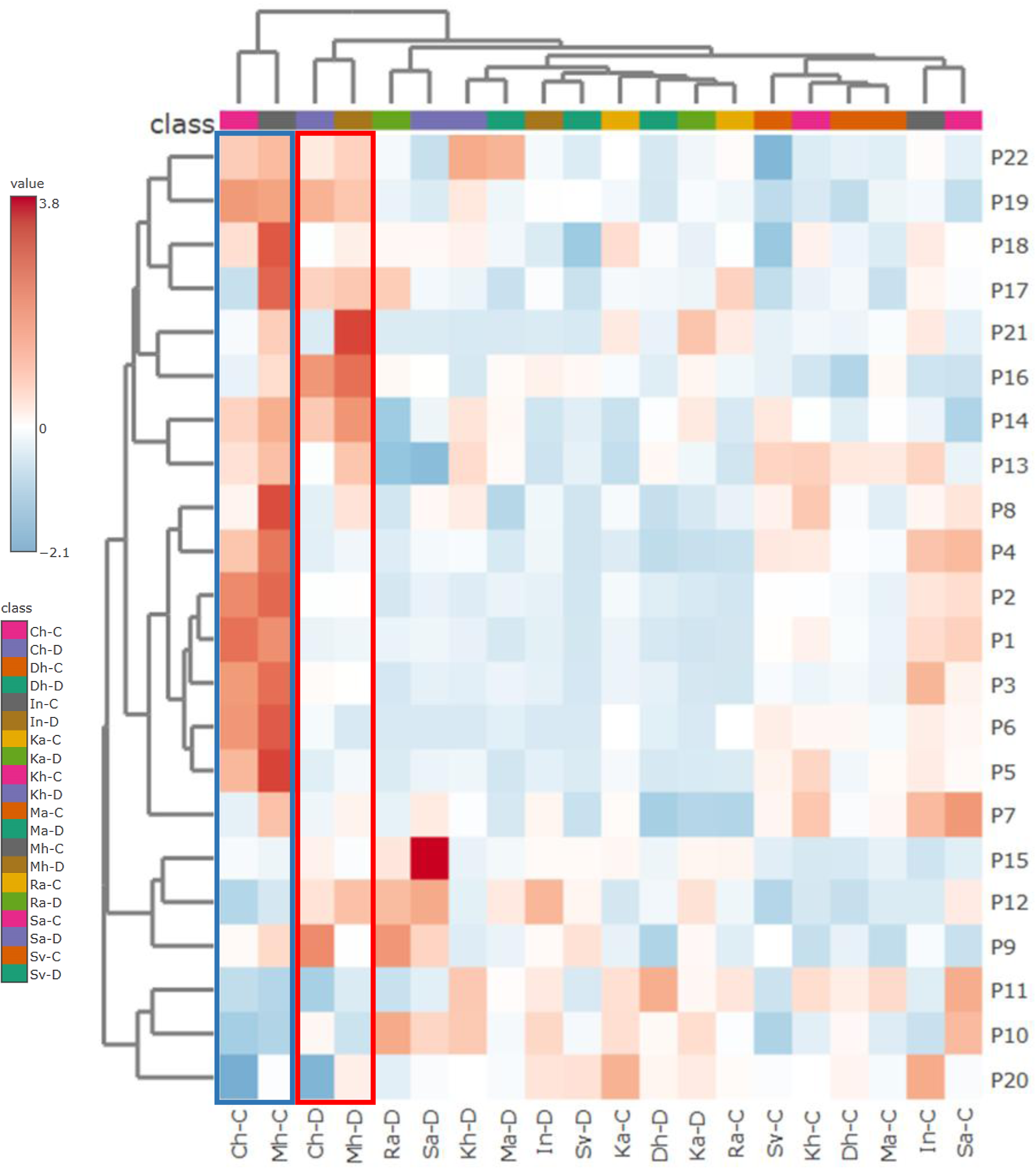
Hierarchical clustering heatmap of the ten genotypes under control and drought stress based 22 traits and grown in rhizotron. P1: Weight of Stem(s), leaves (g); P2: Weight of Ears (g); P3: Weight of the Plant; P4: Weight of the Grains (g); P5: Quantity of all grains/plant; P6: Quantity of Ears/plant; P7: Weight of the grains of the main ears; P8: Quantity of grains of the main ears; P9: N content in soluble fraction; P10: C content in soluble fraction; P11: Ratio C/N in soluble fraction; P12: δ¹³C in soluble fraction, P13: N content in solid fraction; P14: C content in solid fraction; P15: Ratio C/N in solid fraction; P16: δ¹³C in solid fraction, P17: Total Root Length BBCH 30; P18: Total Root Length BBCH 60; P19: Total Root Length BBCH 75; P20: Root Tips BBCH 30; P21: Root Tips BBCH 60; P22: Root Tips BBCH 75. The heatmap was constructed using Metaboanalyst with default settings. Phenological stages: BBCH 30: Stem elongation; BBCH 60: onset of flowering; BBCH 75: grain development. Abbrevations: Ch: Chili; Mh: Mahmoudi; Dh: Dhahbi; IN: INRAT100; Ka: Karim, Kh: Khiar; Ma: Maali; Ra: Razzek; Sa: Salim; Sv: Svevo.

### Durum Wheat Varieties develop distinct transcriptional programs

To understand the transcriptional programs underlying the phenotypic plasticity observed in the 10 durum wheat varieties, grown in rhizotrons under water stress, we performed a transcriptomic analysis of the flag leaves, whose development is crucial for grain filling. We considered differentially expressed genes (DEGs) as those having |Log2FC| > 1 with a FDR < 0.05. The number of significant DEGs varied among genotypes, with the highest observed in Karim and Khiar (4794 and 4271, respectively) whereas Razzek and Dhahbi had only 334 and 133 DEGs, respectively (Fig. 6A). The clustering heatmap combining all the DEGs illustrates well the different expression profiles, suggesting that distinct transcriptional programs might take place in the 10 genotypes in response to drought stress (Fig. 6B). This assumption was reinforced through an UpSet plot generated to distinguish the common from genotype-specific DEGs across the 10 genotypes (Fig. 6C). While a small number of common DEGs were shared by a subset of genotypes, many appeared to be genotype-specific DEGs. In details, the highest numbers of unique DEGs were found in Karim and in Khiar (2614 and 2279) followed by Maali (1136). Razzek, Svevo and Dhahbi displayed the fewest unique DEGs (20, 37 and 8, respectively), while the two landraces Chili and Mahmoudi exhibited 607 and 91 unique DEGs, respectively. No DEG was common to the 10 genotypes and only 117 DEGs overlapped between different combinations of 4 genotypes (Fig. 6C). Using Mercator, we could identify the number of DEGs enriched across different biological processes in the different genotypes (Fig. 6C, insert). These pie charts showed that processes like photosynthesis, solute transport, protein homeostasis are differentially affected by drought in the different cultivars and noteworthy that cultivars like Svevo and Chili mostly up-regulate genes whereas Karim turned off most of its transcriptional program.

**Figure 6:**
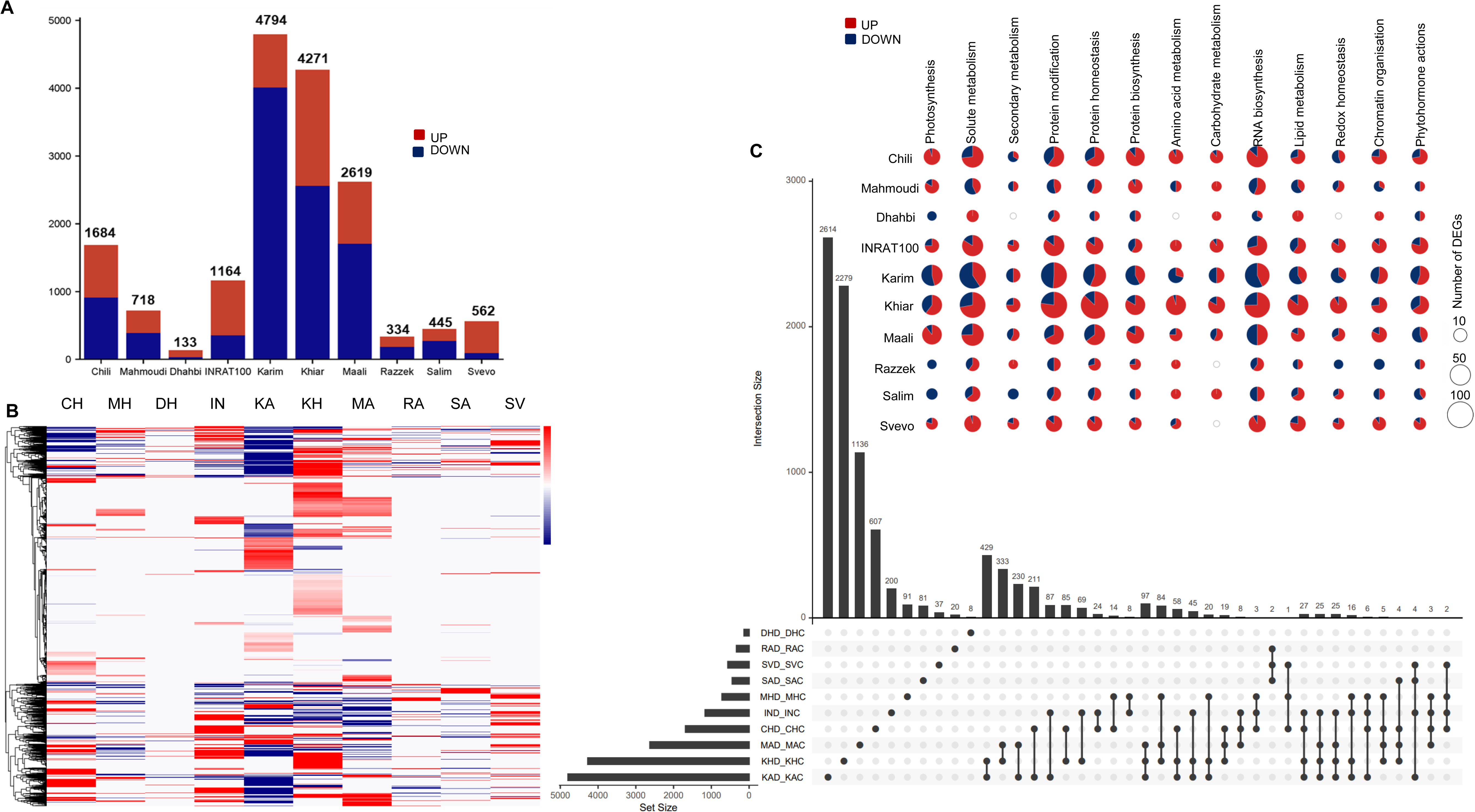
(A) Number of DEGs found in the different genotypes (Chili (CH), INRAT100 (IN), Karim (KA), Khiar (KH), Maali (MA), Mahmoudi (MH), Razzek (RA), Salim (SA), Svevo (SV)) with a cutoff of |log2FC| >1 and FDR<0.05. (B) Clustering heatmap of all the DEGs with a cutoff of |log2FC| >1 and FDR<0.05 in the ten genotypes. (C) UpSet plot showing the size and composition of intersections among DEGs detected in the durum wheat genotypes. The bar chart (top) indicates the number of DEGs in each intersection; filled dots connected by lines (bottom) denote the genotypes contributing to a given intersection, while grey dots indicate absence. The bar chart on the left side indicates the number of DEGs in each genotype. The insert represents pie charts of the number of DEGs for each genotype and for different biological processes extracted using Mercator4 v8.

Because photosynthesis is one of the first processes negatively affected by drought stress, we conducted a comparative analysis of DEGs related to this key biological pathway. Consistently there was a strong down-regulation (net counts = down vs up) of core photosynthesis genes in all genotypes but with a particular genotype-specific responses. Khiar exhibited the highest number of DEGs related to photosynthesis (117) among which 91 were down-regulated and 26 up-regulated (Fig. 7). Khiar and Karim presented a strong repression of genes building photosytem II (antenna (LHC-b) and Calvin cycle genes important for photoreactions and carbon fixation. For instance, Khiar had 13 DEGs encoding LHCb component which were all down-regulated and a unique down-regulated DEG involved in Cytochrome b6 maturation (Fig. 7; Supplementary Table S1). Moreover, there was a differential transcription of ATP synthase genes and photoprotection-related transcription. While Khiar, Karim and INRAT100 strongly down-regulated photoprotection-related genes, other genotypes like Chili, Mahmoudi and Svevo strongly up-regulated these genes indicative of their switch towards photosynthesis protection and resilience. We noticed 3 and 7 DEGs, which were unique for Mahmoudi and Chili, respectively. Among those, two genes were significantly up-regulated, one encoding the CF0 subunit c ATP synthase (+25,7) in Mahmoudi, and another encoding NAD(P)H-quinone oxidoreductase subunit I (+24,8) in Chili. Both are crucial components for photosynthesis (Rochaix, 2011; Ma *et al*., 2021; Wang *et al*., 2025) and their up-regulation in landraces indicates that they protect photosynthesis and regulate the ATP stocks under drought. Interesting was the differential expression of *ELIP* (Early Light-Induced Proteins) genes which are required for PSII repair and reassembly to protect the photosystem from photoinhibition during high light but also during dessication (Hutin *et al*., 2003). We noticed a strong genotype-dependent regulation of *ELIPs*. Cultivars like Karim and Maali had respectively 11 and 4 *ELIP*s down-regulated with large fold change (−24 in Maali, Supplementary Table S1). In contrast, three *ELIPs* were up-regulated (+22, +24.3 and +5 Log2FC) in INRAT100, while Khiar exhibited a balanced transcriptional regulation of *ELIPs* with 2 genes being up-regulated (+22 and +8) and one strongly repressed (−21). For Mahmoudi and Chili, there was no significant differential expression of *ELIP*s suggesting that these sentinels seem not to play a critical role in protecting photosynthesis in these 2 landraces. RNA-Seq data also showed a net down-regulation of genes related to photorespiration like glycolate oxidase, hydroxypyruvate reductase or glycine decarboxylase. Maali, Khiar and Karim showed significant down-regulated DEGs involved in photorespiration: 5 DEGs in Maali (4 down and 1 up-regulated; 3 DEGs in Karim (all down-regulated) and 4 DEGs in Karim. Noteworthy, glycolate oxidase was strongly up-regulated in Khiar (+26) meaning that this genotype acted to protect photosynthesis by recycling photosynthesis byproducts but glycolate oxidase was down-regulated in Karim (−22) indicating that this genotype rather repressed photorespiration. Chili exhibited down-regulation of glycolate oxidase (−9) while up-regulated glycine decarboxylase (+1,3) suggesting the maintenance of photorespiration. These data indicate that the different accessions regulate differently their photorespiration components in response to drought and that their transcriptional control may depend on the timing and genotype-specific response.

**Figure 7:**
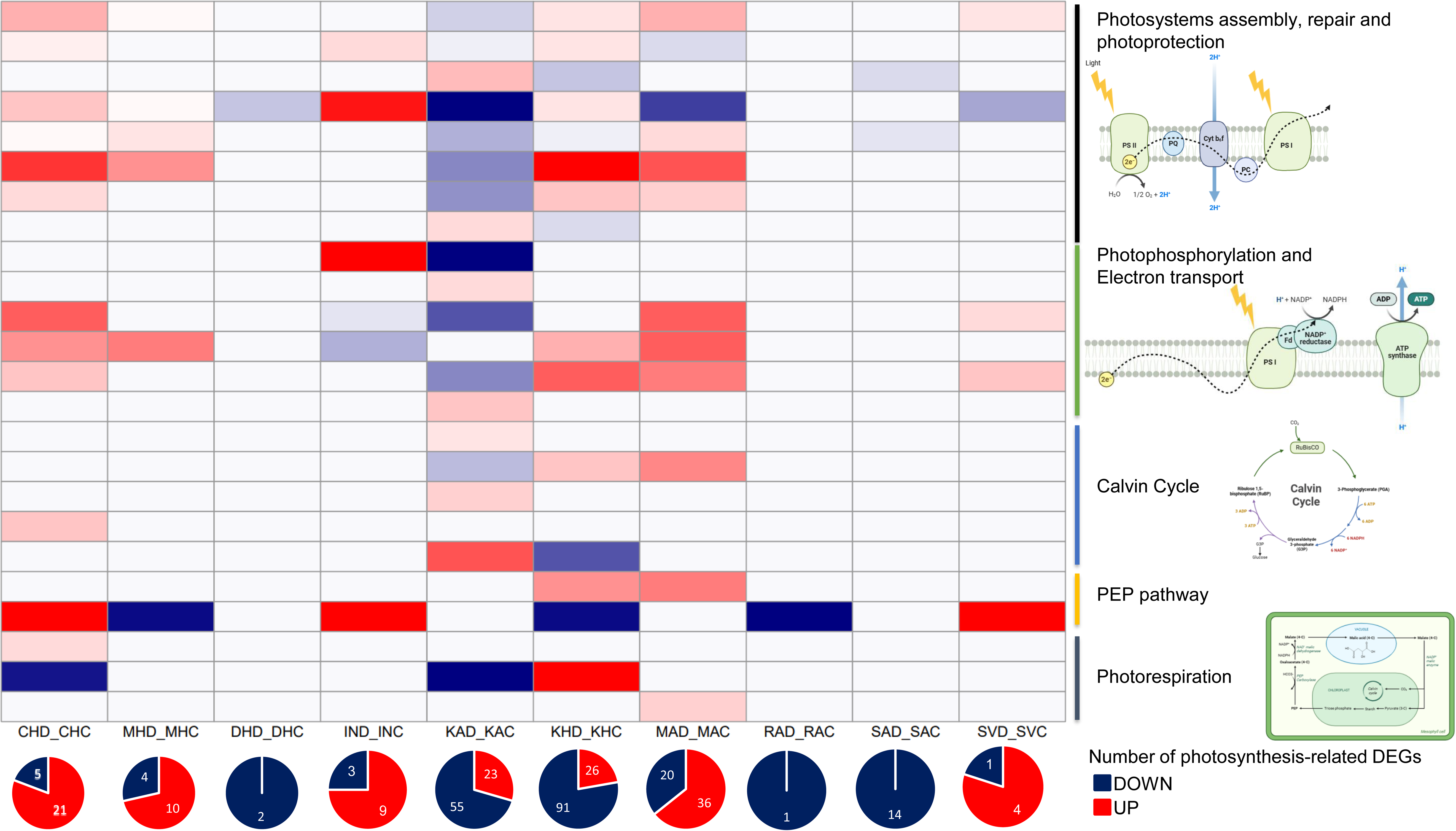
The ten durum wheat genotypes differentially regulate photosynthesis processes. Heatmap showing DEGs expression (defined as |log2FC| >1; FDR<0.05) in different photosynthesis processes. The DEGs were extracted using the bin classification from Mercator4 v8 for each process. The pie charts below represent the total number of up-(red) and down-regulated (dark blue) genes in each genotype. Sketch of the different photosynthesis components were represented using BioRender.

Next, we noticed a pronounced genotype-specific enrichment in DEGs related to solute transport, in all genotypes, except Mahmoudi. Genes involved in nutrient distribution along the plant were strongly down-regulated in Karim, Khiar and Maali but over-expressed in Svevo and INRAT100, suggesting active osmotic/ionic adjustment in those two genotypes (Supplementary Fig. S10).

Several DEGs that are involved in stomatal movement were also identified and remarkably, *TRITD3Av1G264100* encoding an ABA-importer (ABCG40) is up-regulated in Chili, contrasting with its down-regulation by drought in Mahmoudi like its homeologue (−18.89 and −19.22, respectively). Concomitantly, an ABA-exporter (*TRITD4Av1G171070*) was also up-regulated about 4 times in Chili. Other DEGs that are known to play relevant role in drought tolerance were identified in Chili, such as the genes encoding PIP1;4 water channels and the vacuolar transmembrane H^+^-pyrophosphatase. These genes appeared unchanged in Mahmoudi reinforcing the fact that the two landraces adopt two distinct transcriptional programs to drive stress resilience.

Seventy-four transcription factors were de-regulated within the DEGs of all genotypes covering the different structural classes of transcriptional regulators (i.e. bZIP, MYBR2 R3, WRKY; Supplementary Fig. S11; Supplementary Table S1). Notably, Karim had the highest number of down-regulated transcription factors (23 down vs 5 up). In Chili, among the 16 TFs (11 up vs 5 down), the *Argonaute 1* (*AGO1*) homolog and the *Nuclear Factor YA* which were up-regulated. We also noted nine WRKYs in this landrace, among which three were repressed and six were up-regulated. Interestingly, all the seven TFs found in Mahmoudi were down-regulated and belong to WRKY (WRKY28/45/50/76), LUX/BOA (2) and TCP (*TEOSINTE BRANCHED1/CYCLOIDEA/PROLIFERATING CELL FACTOR1 TCP20*) families (Supplementary Fig. S11).

Overall, these RNA-seq data reveal that these shifts in gene expression align with what is often observed under drought: down-regulation of light harvesting/Calvin cycle and up-regulation of membrane/extracellular defense, transport, and stress-responsive TFs, with the magnitude and direction being genotype-dependent. While Karim, drought sensitive, strongly repressed drought-related gene expression like those involved in photosynthesis and solute transport, other genotypes (Chili and Svevo) rather over-expressed these genes.

## Discussion

Durum wheat is a cornerstone crop of the Mediterranean basin, where its production is increasingly constrained by climate change–associated stresses such as rising temperatures and recurrent drought. Mediterranean durum wheat landraces, shaped by centuries of selection under marginal, water-limited conditions, harbor a rich adaptive genetic diversity for stress resilience, phenological plasticity, and resource-use efficiency. However, this unique source of numerous adaptive alleles remains largely underexploited in contemporary breeding programs. The integration of multi-omics approaches enables the systematic characterization of this untapped diversity, facilitating the identification of stress-signaling pathways and the introgression of robust adaptive traits into elite breeding material. This study aimed to harness the genetic diversity of a subset of Tunisian durum wheat varieties in response to drought stress, by using an integrated approach combining HTP, physiological and transcriptomic analyses. We included in our analyses, two Tunisian landraces (Chili and Mahmoudi) seven modern breeding lines alongside the reference variety Svevo. Mahmoudi and Chili are among the most emblematic ancient durum wheat varieties in Tunisia (Ouaja *et al*., 2021). They are well appreciated for their remarkable grain quality, genetic value, and strong agro-ecological adaptation but today they are cultivated by a limited number of traditional farmers.

HTP was performed on this core collection of durum wheat genotypes grown up to maturity under well-watered or long-term drought conditions in pots and rhizotrons, to assess growth, yield components, root architecture, physiological traits, and reaction norm plasticity. The measurement of morphological, agronomic and physiological traits revealed that the two landraces Mahmoudi and Chili performed better than the elite varieties under drought. For instance, the yield parameters including grain quantity/plant, weight of ears, weight of grains are higher in the two landraces under drought stress compared to the other genotypes. It is worth noting that although drought stress caused a stronger relative reduction in yield components in the two landraces, their absolute values remained consistently higher than those observed in modern varieties. This indicates that landraces seem to retain a superior yield capacity under water-limited conditions, highlighting their intrinsic robustness and adaptive potential in drought-prone environments. Similar observations were previously reported by Frankin *et al.,* (2021), where in-field study revealed that several wheat landraces showed good agronomic potential across environments and outperformed modern varieties under drought stress. Our assumption was supported by a series of stress tolerance indices reflecting productivity (MP, GMP, STI) and stability (TOL, YSI, SSI). Based on these indices, the two landraces ranked highest in overall performance, with Mahmoudi showing a clear balance between productivity and stability under drought stress. This trend was clearly illustrated in the PCA biplot, where the two landraces clustered closely together and were followed by INRAT100, while the remaining genotypes appeared more distant.

In addition to the above-ground traits, the analysis of belowground growth provided further insights into the mechanisms adopted by the wheat accessions when facing water stress. Measurements of key root traits, such as total root length, root surface and number of root tips, revealed again a genotype specific response to drought across the three developmental stages (BBCH 30, BBCH 60, and BBCH 75). For instance, despite showing a reduction in total root length under drought, Chili and Mahmoudi appeared to maintain the most robust root system architecture across drought panels, successfully preserving a large root surface area and high numbers of root tips. Both landraces adopted similar strategies, though with subtle distinctions. Notably, at BBCH 60, Mahmoudi exhibited a strong increase in root tips number, compared to Chili. This phenotypic pattern in Mahmoudi suggests a shift in developmental strategy from elongation to increased branching, thereby enhancing root surface area and improving topsoil foraging efficiency as well as nutrient uptake potential. At BBCH 60, a vegetative to early reproductive phase, during which adequate water availability is essential for successful pollination and grain set, Mahmoudi appears to avoid investing in deeper soil penetration to conserve carbon and energy. Such plastic responses of wheat root systems under water deficit have been widely reported and are considered as important mechanisms for maintaining water uptake under drought conditions (Lynch, 2018; Li *et al*., 2021). Therefore, exploring the genetic loci regulating root traits in this landrace could provide valuable molecular targets for breeding wheat with enhanced drought tolerance.

On the other hand, we noticed that yield and growth performance of the two landraces under drought correlated well with several stress tolerance indices and physiological traits such as iWUE, C and N partitioning. Leaf insoluble C and N partitioning values are reliable indicators of the accumulation of structural components, including cell wall–associated carbon (e.g. lignin and cellulose) and nitrogen associated with photosynthetic proteins. These values were consistently higher in the two landraces compared with the other genotypes, suggesting a strategy oriented towards long-term survival and biomass preservation under water-deficit conditions explaining their yield robustness under drought (Le Gall *et al*., 2015). Moreover, we observed a relatively more balanced C/N ratios in leaves of Mahmoudi and Chili, compared to other varieties, suggesting that they may maintain sufficient photosynthetic carbon supply relative to nitrogen assimilation, which supports metabolic activity under drought and may facilitate a faster recovery once water availability is restored. This assumption is in line with previous findings showing that drought can lead to imbalanced leaf C/N ratios that are linked with photosynthetic decline and stress-induced physiological responses as it was reported in sorghum and bread wheat (Chen *et al*., 2015; Shokat *et al*., 2024).

To underpin the differential physiological behavior of the two landraces, we conducted a comparative analysis of leaf iWUE and Ψπ. Once more, the two landraces surpass the breeding lines and exhibited the highest iWUE values in the leaf solid fraction. This pattern is indicative of a more efficient regulation of carbon assimilation relative to water loss under drought conditions, likely reflecting tighter stomatal control (Condon *et al*., 2004; Lawson and Blatt, 2014). Such physiological adjustment further supports improved photosynthetic efficiency under water stress and is consistent with the higher drought tolerance observed in these landraces. In the case of the soluble fraction, the shifts in the iWUE observed under drought across all the genotypes (Supplementary Fig. S8) most likely reflect short-term (hours or days) changes in stomatal conductance and carbon assimilation and allocation. These rapid changes may be associated with metabolic reprogramming under drought, potentially affecting the allocation of carbon to osmolyte biosynthesis and compatible solute production to maintain cellular function under water stress (Farquhar *et al*., 1989; Mininni *et al*., 2022). The two landraces, Chili and Mahmoudi, demonstrated solid-fraction stability and adaptive soluble-fraction responses which may reflect a sophisticated drought adaptation approach that balances structural maintenance with metabolic flexibility. In line with these acute physiological adjustments, the two landraces, especially Chili exhibited the most negative Ψπ values suggesting its higher capacity for osmotic adjustment under drought, probably via the accumulation of osmo-protective compounds. Therefore, we can deduce despite some phenotypic similarities as shown by the clustered heat map, that the two landraces seem to deploy distinct physiological mechanisms especially those controlling the short-term response to drought.

Based on these various phenotypical and physiological analyses, it becomes clear that the two landraces showed a better resilience to drought, compared to the modern durum wheat varieties. Moreover, they show also a more pronounced phenotypic plasticity to water stress. It should be noted that drought tolerance and phenotypic plasticity are not always closely correlated as phenotypic plasticity is not necessarily adaptive (Nicotra *et al*., 2010; Tardieu, 2012). Phenotypic plasticity contributes to drought tolerance when trait adjustments are adaptive, as illustrated by the rapid iWUE shifts observed in the leaf soluble fractions of the two landraces. In such cases, plasticity helps buffer performance losses, allowing yield and/or biomass to remain relatively high. Conversely, strong plastic responses may also reflect stress sensitivity, particularly when they are associated with pronounced reductions in growth, photosynthesis, or yield.

By integrating the phenotypical and physiological traits with transcriptomic data we aimed at dissecting the molecular mechanisms behind the drought tolerance and phenotypic plasticity of these two landraces. The choice of the flag leaf is not fortuitus as it contributes for up to 50–60% of the carbon assimilates that fill the grain and under drought the photosynthetic performance of the flag leaf largely determines grain yield and quality (Biswal and Kohli, 2013). Therefore, such transcriptomic analyses can directly link gene expression changes to yield stability and drought tolerance.

The transcriptomic analysis indicated substantial variation in the number of significant DEGs among genotypes with some cultivars showing strong transcriptional reprogramming (Karim and Khiar) whereas others exhibiting weak responses (Dhahbi, Razzek, Salim). These patterns indicate that the magnitude of transcriptional change is not proportional to the degree of drought tolerance. This is also noteworthy in the two landraces with Chili modifying the expression of twice more genes than Mahmoudi. However, the literature shows both patterns: in some studies, stress-sensitive genotypes show massive transcriptional disruption, (Kumar et al., 2018; Lv et al., 2022) when others, tolerant genotypes display higher numbers of DEGs. (Lv *et al*., 2018; Chaichi *et al*., 2019). In our case, the two landraces seem to exhibit pre-adapted or “constitutive” drought-tolerant expression states, while some sensitive genotypes (i.e Karim) need a larger emergency response, hence more DEGs, consistent with the “transcriptional shock vs. transcriptional poise” framework for plant stress responses (Harb and Samarah, 2015; Khan et al., 2025). Accordingly, the drought tolerance observed in the current study is not driven by the number of DEGs but rather depends on the functional relevance and coordination of specific regulatory and physiological pathways such as photosynthesis. Among the photosynthesis-related genes that are affected by drought in different breeding lines, we noticed those encoding *ELIPs*, chloroplast proteins known to play an essential role to protect photosynthesis from high-light-induced damage (Hutin *et al.,* 2003). *ELIPs* have been reported to be induced not only by highlight but also by other environmental stresses, including drought (Bartels *et al.,* 1992). In our transcriptomic study, many *ELIP* genes showed very large foldchanges, but in opposite directions across genotypes (some massively up, others down), suggesting different acclimation strategies among the ten varieties. Interestingly, the transcription of these *ELIP*s was not significantly affected by drought in the two landraces, indicating that they might be not relevant in this stress response or are part of the pre-adaptive responsive genes. However, genes involved in photophosphorylation and electron transport were mainly up-regulated in the two landraces, suggesting their ability to sustain energy production and photosynthesis during stress. ATP synthase was up-regulated in Mahmoudi, and the oxidoreductase subunit was up-regulated in Chili. Both are crucial components for photosynthesis, especially under drought where plants face low CO_2_ because of stomal closure. The CF0 subunit c of ATP synthase forms part of the proton channel in the thylakoid membrane, which drives ATP synthesis, prevents proton overload, stabilizes NPQ, and maintains safe electron flow, keeping the photosynthetic machinery alive when CO₂ becomes limiting because of stomatal closure under drought (Wang *et al.,* 2025). Moreover, NAD(P)H-quinone oxidoreductase subunit I (NDH-I), part of the NDH complex, that enables NDH-dependent cyclic electron flow to generate ATP, protects photosystem I from over-reduction, maintains proton balance, limits ROS, and sustains photosynthesis when CO₂ is scarce (Ma *et al.,* 2021). Although functionally different, NDH-I and CF0 play complementary roles as both influence ATP supply, proton motive and photoprotection force under drought, suggesting that they might be part of the same adaptive system that wheat cultivars use during water stress. Beyond these genes, we noticed that drought has resulted in a net down-regulation (counts down vs up) of other genes involved in photosynthesis-related pathways, particularly Cytochrome b6_66/f (9 vs 1), Calvin cycle (25 vs 6), Photorespiration (11 vs 3), PSII LHC (antenna; 20 down vs 6 up-regulated). This down-regulation of the core photosynthesis mirrors other wheat transcriptomes reporting broad suppression of photosystems and carbon fixation during dehydration (Zuluaga *et al.,* 2023; Cevher-Keskin *et al.,* 2025).

On the other hand, our search among the DEGs in the two landraces has resulted in the identification of a small number of genes (2 in Mahmoudi and 14 in Chili) that are well known to be involved in drought response. The two DEGs identified in Mahmoudi are functionally redundant and encode the ABC transporter G family member 40 (ABCG40), mediating the import of ABA in guard cells and controlling stoma closure. Interestingly, these two genes are significantly down-regulated, suggesting that Mahmoudi could adopt a more fine-tuned stomatal closure as an adaptive response to drought. This plasticity in stomatal conductance (with less negative δ¹³C values) may allow continued carbon assimilation without a large water loss, and sustained photosynthesis, which could be advantageous in Mediterranean climates characterized by episodic drought stress.

For Chili, among the 14 drought-responsive genes there are three functionally related encoding glutathione-S-transferases (GST) known to be required for detoxification processes during stress (Marrs, 1996). These genes were down-regulated to different extents, by drought stress. Interestingly, a knockout mutant in the homologous gene of *Arabidopsis thaliana* was reported to be more drought tolerant and accumulate higher levels of glutathione and ABA *(Chen et al.,* 2012). In addition, several DEGs that are seemingly involved in stomatal movement were identified. Remarkably, *TRITD3Av1G264100* encoding an ABA-importer (ABCG40) is up-regulated in Chili, contrasting with its down-regulation by drought in Mahmoudi. Concomitantly, an ABA-exporter (*TRITD4Av1G171070*) was also up-regulated about 4 times in Chili. Other DEGs that are known to play relevant role in drought tolerance were also found, such as the genes encoding PIP1;4 water channels and the vacuolar transmembrane H^+^-pyrophosphatase. Moreover, two regulators were up-regulated in Chili, the Argonaute 1 (*AGO1*) homolog (*TRITD5Av1G209870*) and the Nuclear Factor YA (*TRITD6Bv1G196010*). Several reports showed that the *AGO* genes were up-regulated in several crops by abiotic stress, especially drought (Bai *et al.,* 2012; Zhao *et al.,* 2015). The up-regulation of *AGO1* indicates that an epigenetically regulated drought response may occur in Chili. Nuclear factor YA is the DNA binding subunit of a heterotrimeric transcription factor involved in the transactivation of genes harboring CCAAT cis-elements common for ubiquitous promoters but also those controlling stress responsive genes (Zhao *et al.,* 2017).

Overall, from a mechanistic point of view, the expression patterns in Mahmoudi and Chili are consistent with the superior drought tolerance observed in some durum wheat genotypes, through tighter control of stomatal closure, less negative δ¹³C values, higher iWUE and osmotic adjustment, as well as better regulation of photoprotection and ATP supply in the chloroplasts. The comparative analysis between the two landraces revealed that common and distinct pathways seem to be activated in response to water stress. Both seem to solicit ABA-dependent and ABA-independent mechanisms. In Chili but not Mahmoudi, other pathways relying on transcriptional and epigenetic regulation, reactive oxygen metabolism and detoxification, as well as transport activities might be activated.

Our findings collectively show that drought resilience in durum wheat emerges from the coordinated modulation of physiological plasticity, root system robustness, and targeted transcriptional reprogramming. By integrating whole-plant phenomics with organ-level transcriptomics, this study provides mechanistic clarity on the traits that enable stable performance under chronic drought stress. The contrasting yet effective strategies employed by the landraces Chili and Mahmoudi highlight the value of locally adapted germplasm for uncovering adaptive mechanisms shaped by long-term selection in water-limited environments and may help in accelerating the development of climate-resilient durum wheat cultivars.

## Acknowledgements

The authors thank the gardeners and technical team of the Research Group Environmental Simulation (EUS) and the Research Group of Plant Genome and System Biology (PGSB), at Helmholtz Research Center of Munich, for their experimental and technical assistance. We are also very grateful to Dr. Bassem Bouaziz from MIRACL Laboratory at the Higher Institute of Computer Science and Multimedia, University of Sfax-Tunisia, for his assistance in root image data processing and analysis.

## Author Contribution

CE, MH, JBW, JPS, and KM: conceptualization; JPS, MH, CE and MM: methodology; MM, RD, IG, RT, SC, MG and RB: formal analysis; RD, IG, RT, SC, MG: investigation; CE, MH, RG, AE, KM: resources; MM, JPS, JBW, and CE: data curation; RD, IG, RT, CE and MH: writing - original draft; RD, IG, RT, JPS, CE, KM and MH: writing - review & editing; RD, IG, RT, CE and MM: visualization; CE, JPS, and MH: supervision; CE, RG, AE, JPS, KM and MH: funding acquisition.

## Data availability

Sequencing data have been submitted to EMBL-EBI (https://www.ebi.ac.uk/biostudies) under the ArrayExpress accession number E-MTAB-16860. The raw data of phenotypic and physiological traits are available on Zenodo (https://doi.org/10.5281/zenodo.19392026)

---

**Fig. S1:**
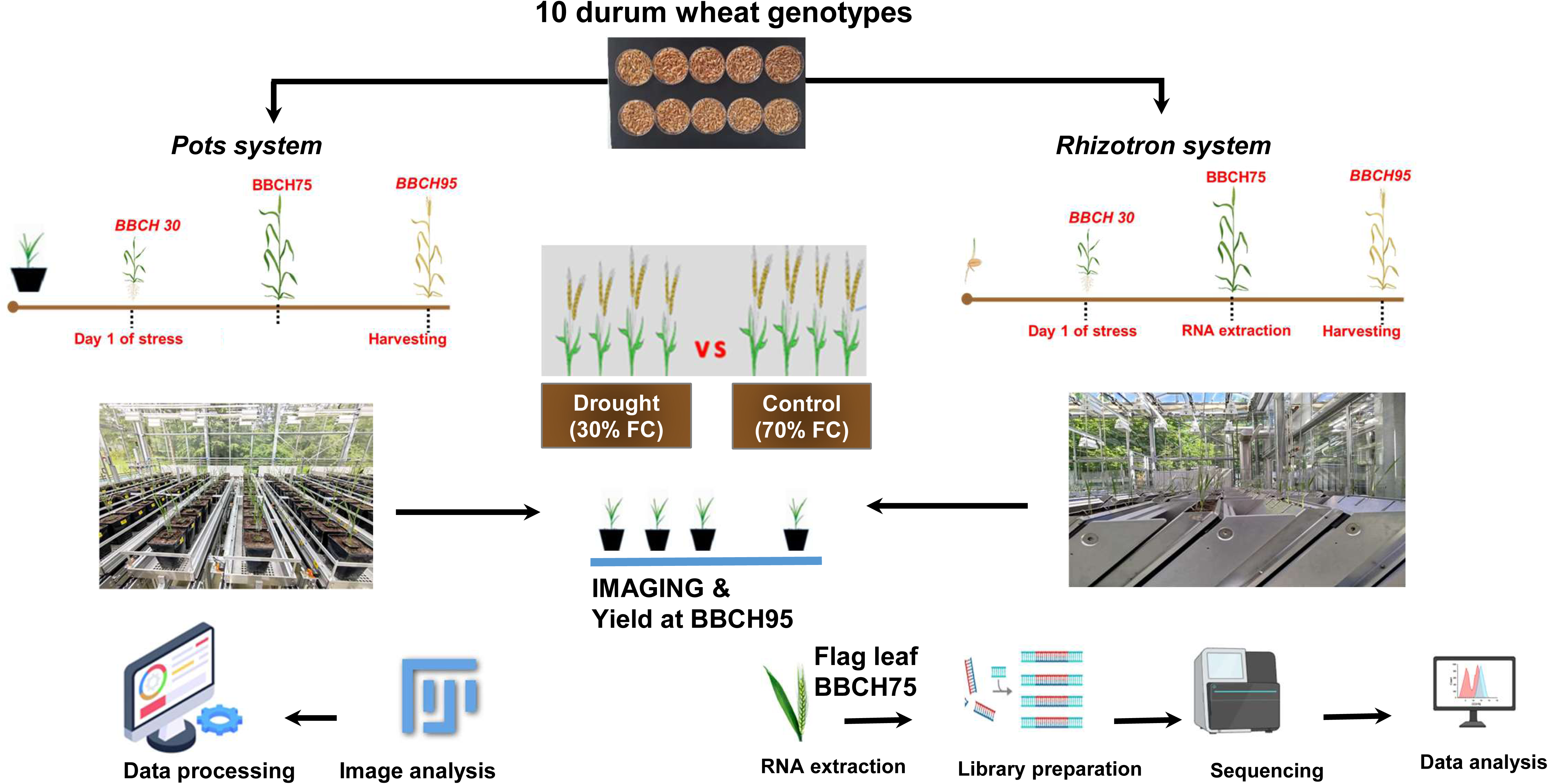
Experimental setup of durum wheat cultivation in rhizotron (R) and pots (P) applied to 10 durum wheat varieties under control conditions (70% nPC) or drought stress (30% nPC) initiated at BBCH 30. Phenological stages: BBCH 30: Stem elongation; BBCH 60: onset of flowering; BBCH 75: grain development.

**Fig. S2:**
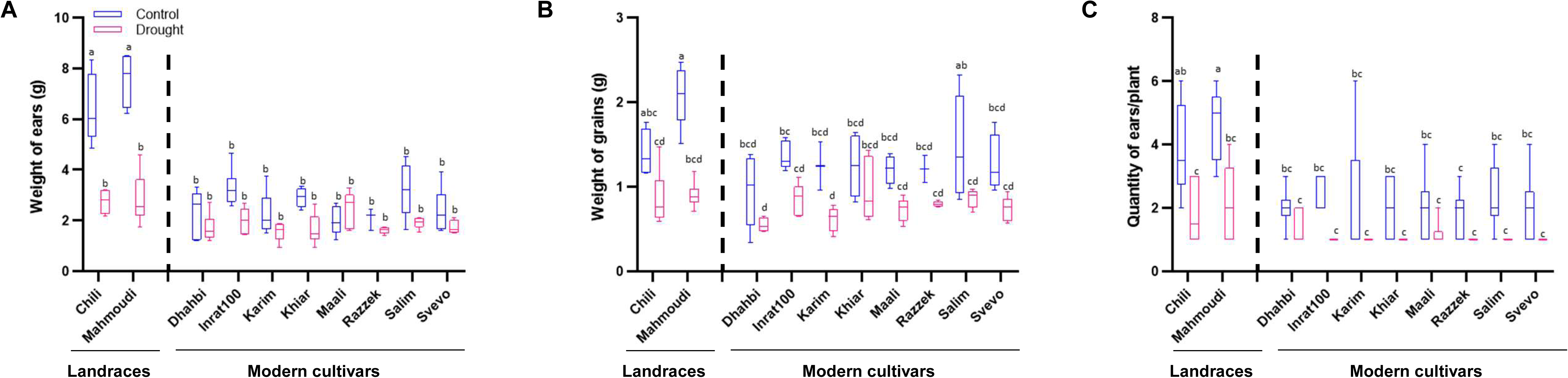
Comparative assessment of yield traits in ten durum wheat varieties under control and drought stress conditions in rhizotron system. (A) Weight of ears (B) Weight of grains. (C) Quantity of ears/plant. Data represent the means and standard deviations (error bars) of six biological replicates for the Rhizotron (R) system. Different letters indicate significant differences (P < 0.05) among samples, as determined by one-way ANOVA followed by Tukey’s HSD test.

**Fig. S3:**
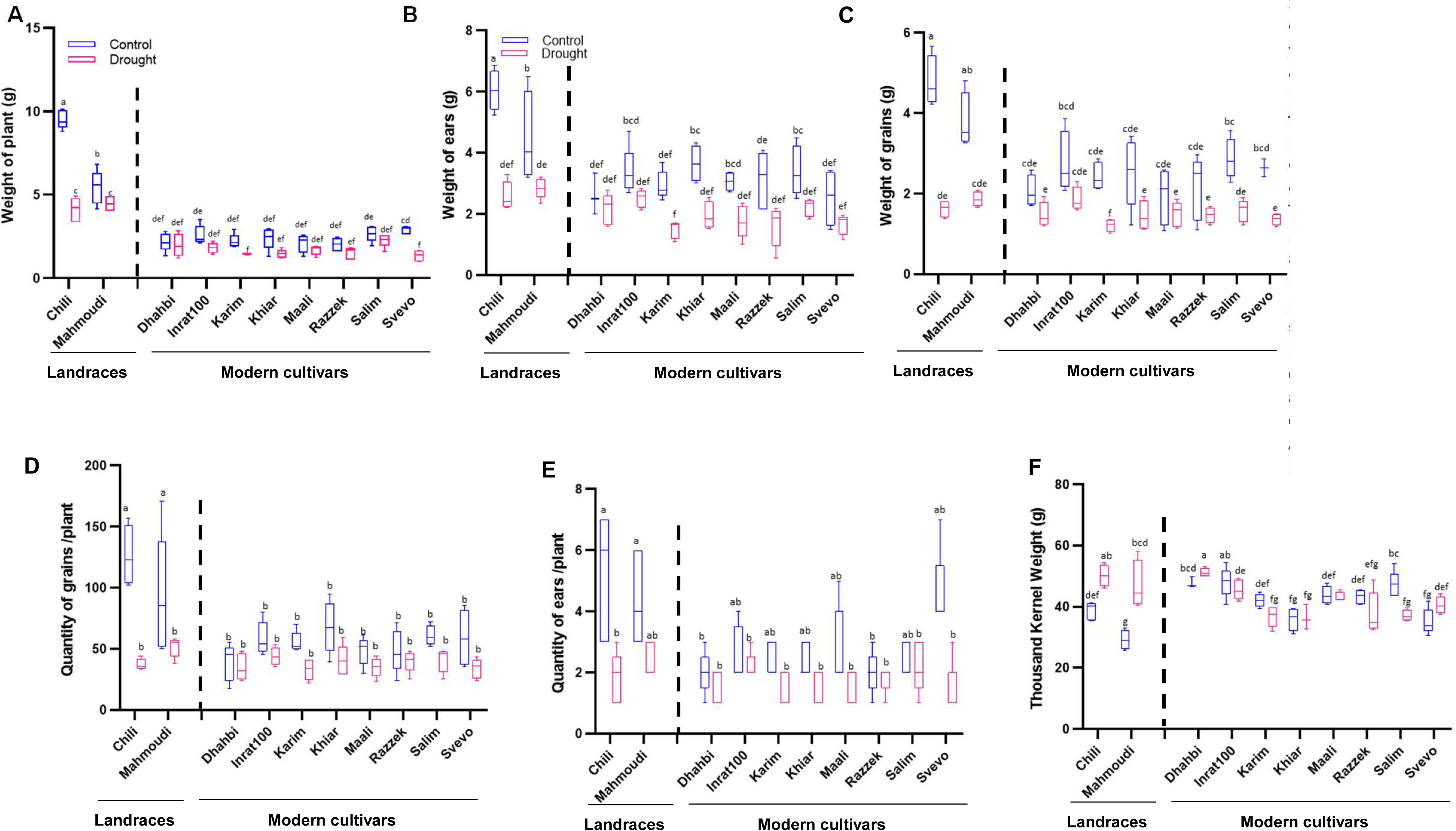
Comparative assessment of yield traits under control and drought stress conditions in pots system. (A) Weight of the plant (B) Weight of ears. (C) Weight of grains (D) Quantity of grains (E) Quantity of ears. (F) Thousand Kernel Weight. Data represent the means and standard deviations (error bars) of six biological replicates for the pot (P) system. Different letters indicate significant differences (P < 0.05) among samples, as determined by one-way ANOVA followed by Tukey’s HSD test.

**Fig. S4:**
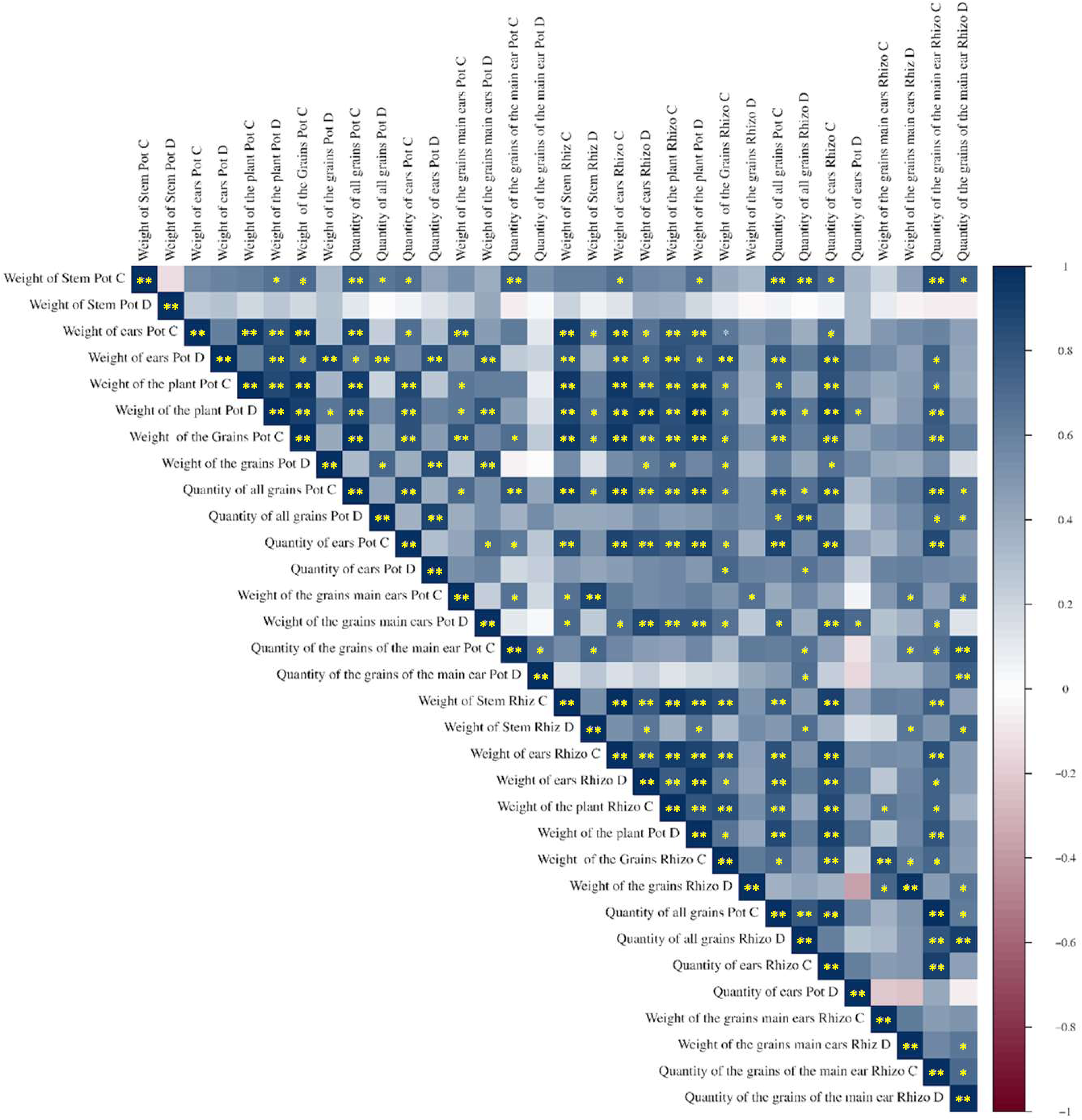
Correlation between drought response traits in durum wheat varieties grown in two systems (pot and rhizotron). Correlations between yield and growth parameters were calculated using data collected under control and drought stress conditions (BBCH 95). Color intensity represents Pearson’s correlation values as shown in the color scale. Significant correlations (P < 0.05) are indicated with an asterisk (*) and (P<0,01) with a double asterisk**

**Fig. S5:**
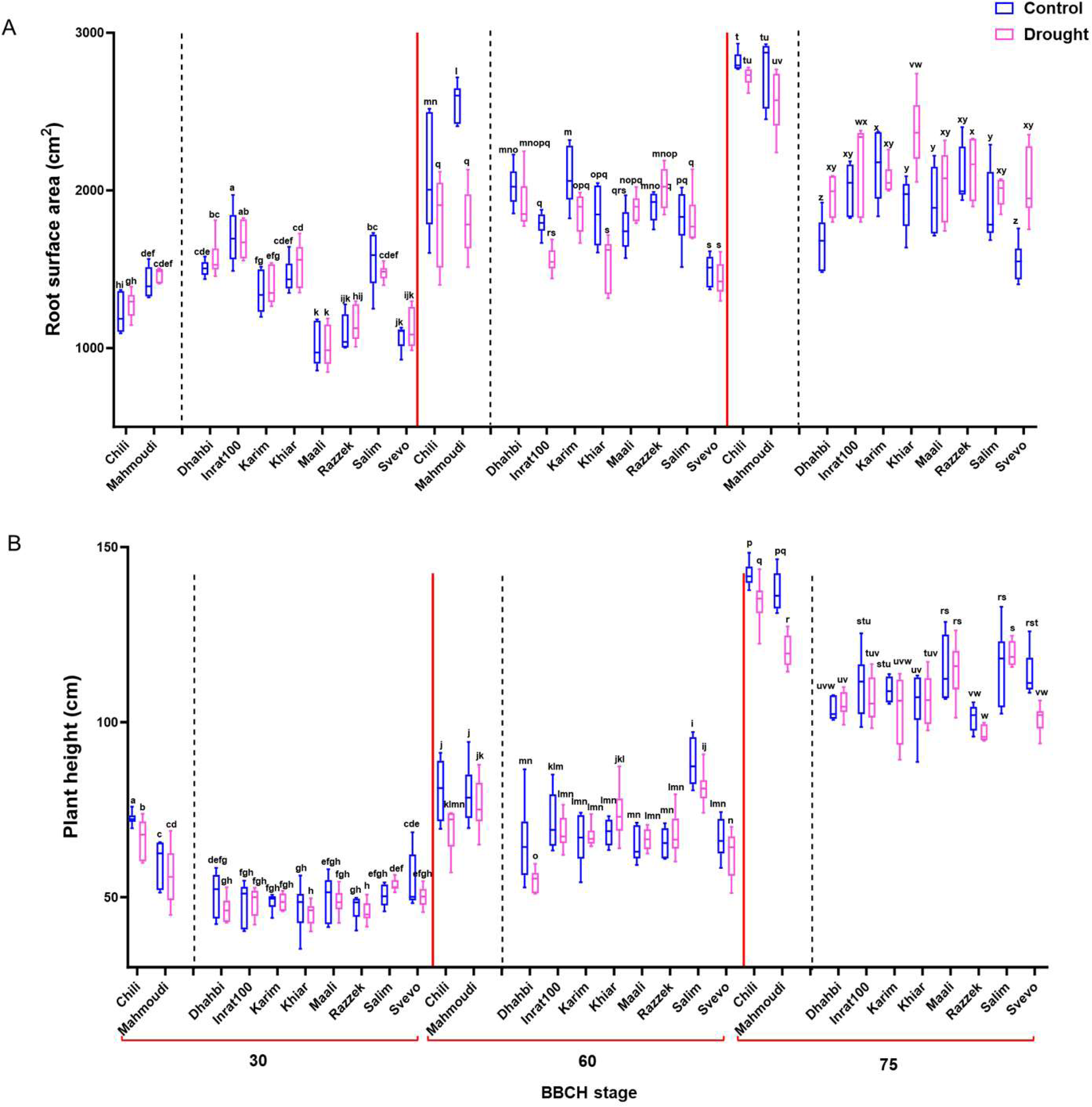
Root surface area (A) and plant height (B) at BBCH 30, BBCH 60, and BBCH 75, grown under control and drought conditions. Root surface area measurements were obtained using RhizoVision Explorer from segmented outputs generated by RootPainter, based on images of roots from plants grown in the rhizotron (n = 6). Plant height was measured using ImageJ from images captured of plants grown in pots (n = 6). Within each panel, treatments of ten genotypes at each BBCH stage were compared using one-way ANOVA followed by Student’s t-test.

**Fig. S6:**
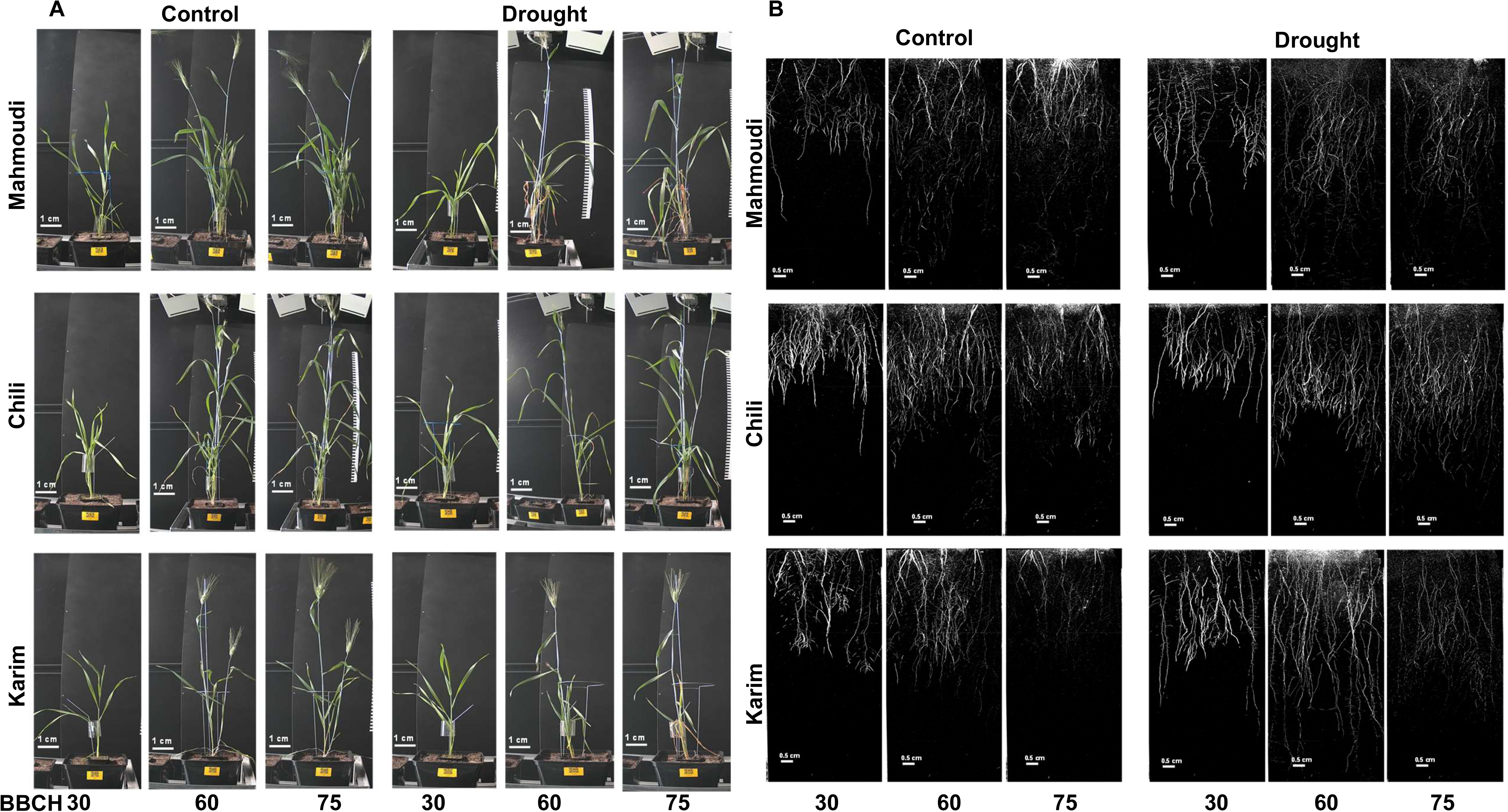
Representative pictures of the plants (A) and their roots (B) of 3 genotypes (Mahmoudi, Chili and Karim) at BBCH30, 60 and 75 grown either under control or drought conditions. Phenological stages: BBCH 30: Stem elongation; BBCH 60: onset of flowering; BBCH 75: grain development. Scale bar (A) 1 cm; (B) 0.5 cm

**Fig. S7:**
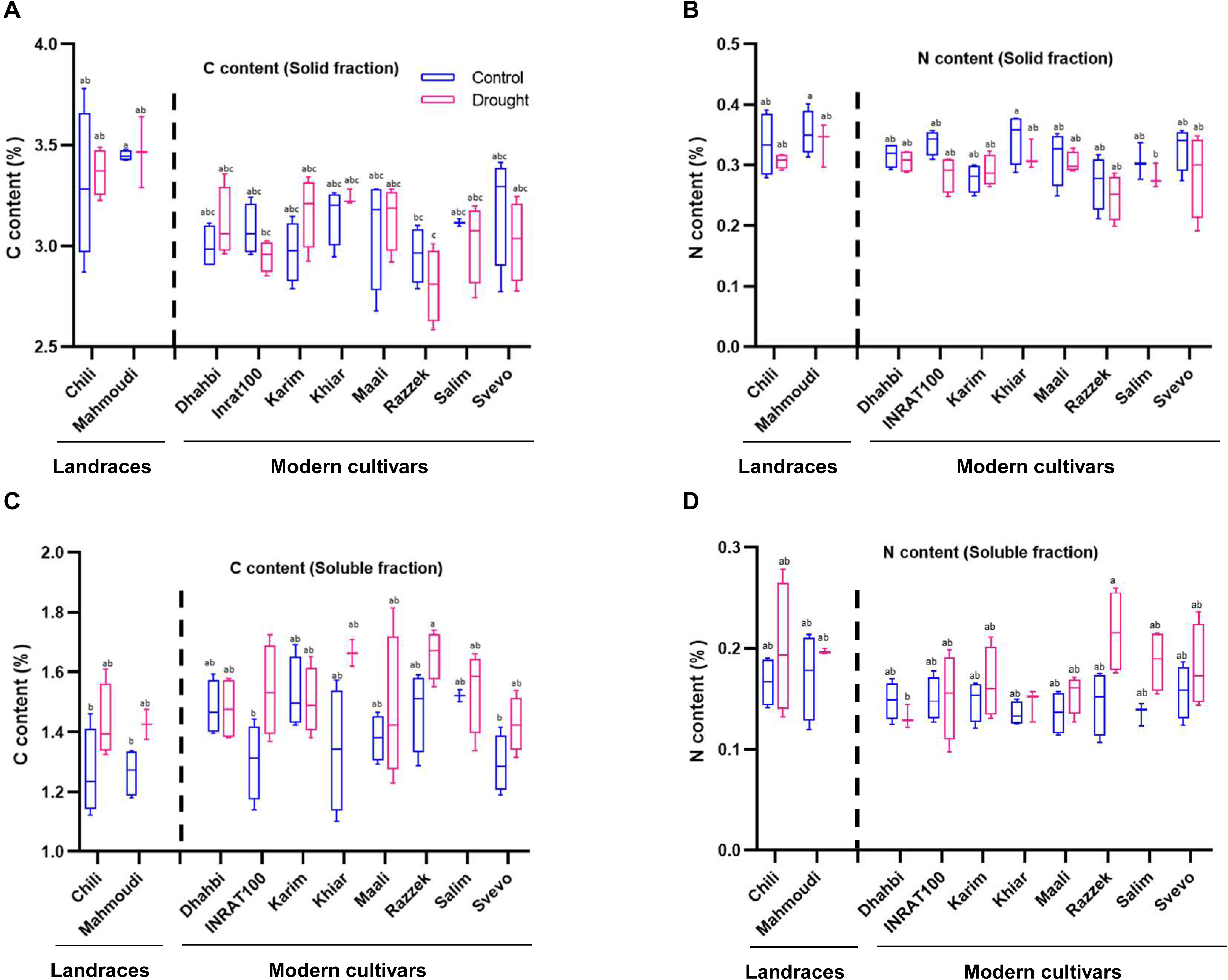
Comparison of Nitrogen (N) and Carbon (C) contents in the leaves of 10 durum wheat genotypes under drought stress conditions. The dynamics of (A, D) carbon content, (B, E) nitrogen content, and (C, F) C/N ratio are shown for both solid and soluble leaf tissue fractions across all varieties. Different letters denote statistically differences between genotypes as determined by one-way ANOVA followed by Tukey’s HSD test (P < 0.05). Data are mean values from four biological replicates.

**Fig. S8:**
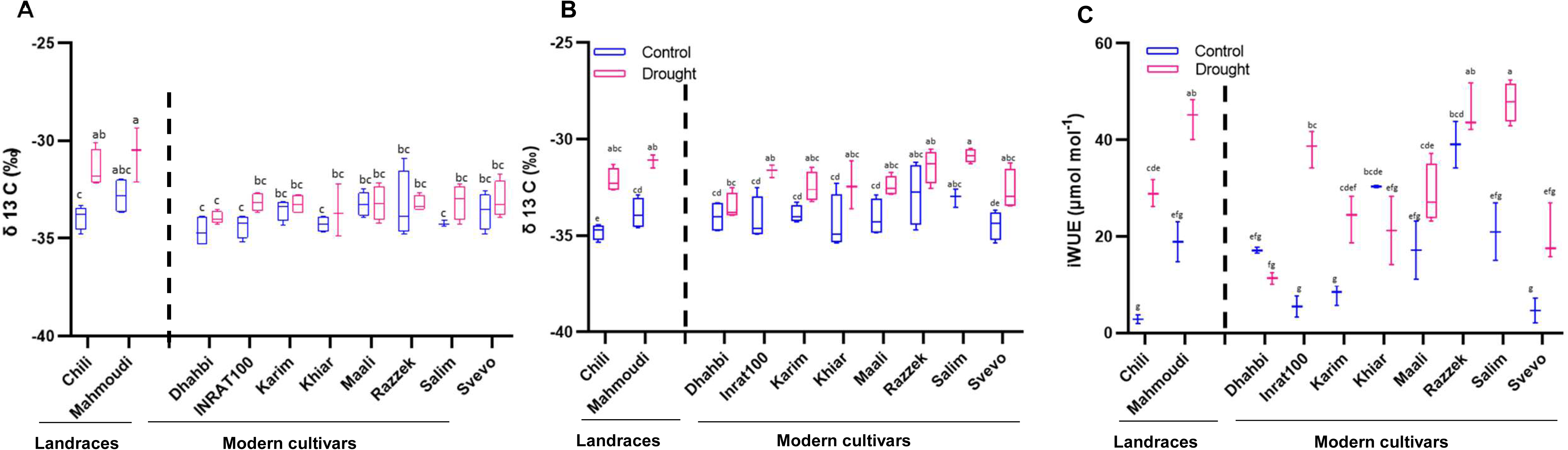
Responses of carbon isotope ratio (δ¹³C) and intrinsic water-use efficiency (iWUE) to drought stress across the ten genotypes (A) Carbon isotype ratio δ¹³C in solid fraction. (B) Carbon isotype ratio **δ¹³C** in soluble fractions (C) iWUE in soluble fractions. Different letters denote statistically differences between genotypes as determined by one-way ANOVA followed by Tukey’s HSD test (P < 0.05). Data are mean values from four biological replicates.

**Fig. S9:**
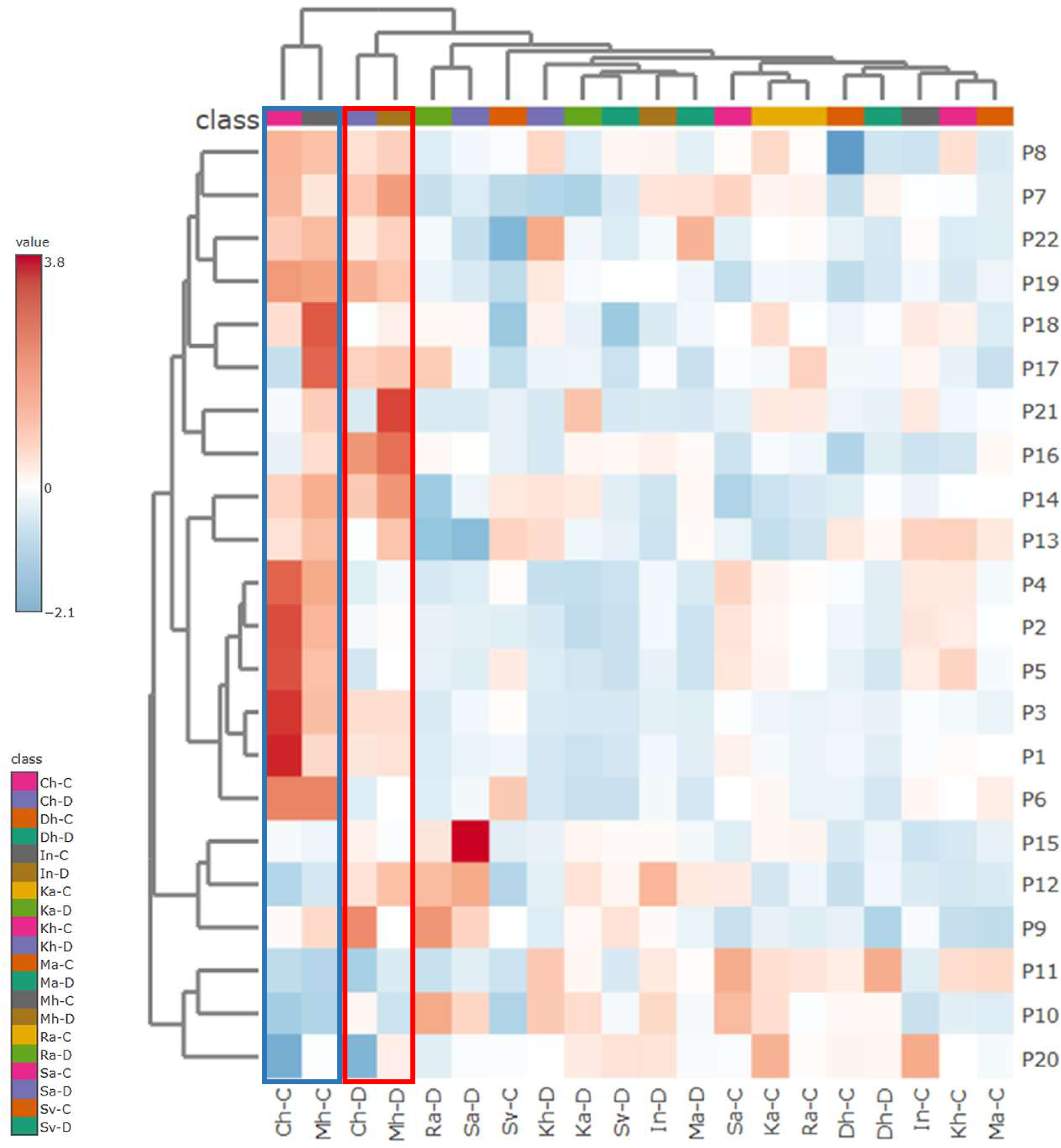
Hierarchical clustering heatmap of the ten genotypes under control and drought stress based 22 traits and grown in pots. P1: Weight of Stem(s), leaves (g); P2: Weight of Ears (g); P3: Dry matter Weight of the Plant; P4: Weight of the Grains (g); P5: Quantity of all grains/plant; P6: Quantity Ears/plant; P7: Weight of the grains of the main ears; P8: Quantity of grains of the main ears; P9: N content in soluble fraction; P10: C content in soluble fraction; P11: Ratio C/N in soluble fraction; P12: δ¹³ C in soluble fraction, P13: N content in solid fraction; P14: C content in solid fraction; P15: Ratio C/N in solid fraction; P16: δ¹³C in solid fraction, P17: Total Root Length BBCH 30; P18: Total Root Length BBCH 60; P19: Total Root Length BBCH 75; P20: Root Tips BBCH 30; P21: Root Tips BBCH 60; P22: Root Tips BBCH 75The heatmap was constructed using Metaboanalyst with default settings.

**Fig. S10:**
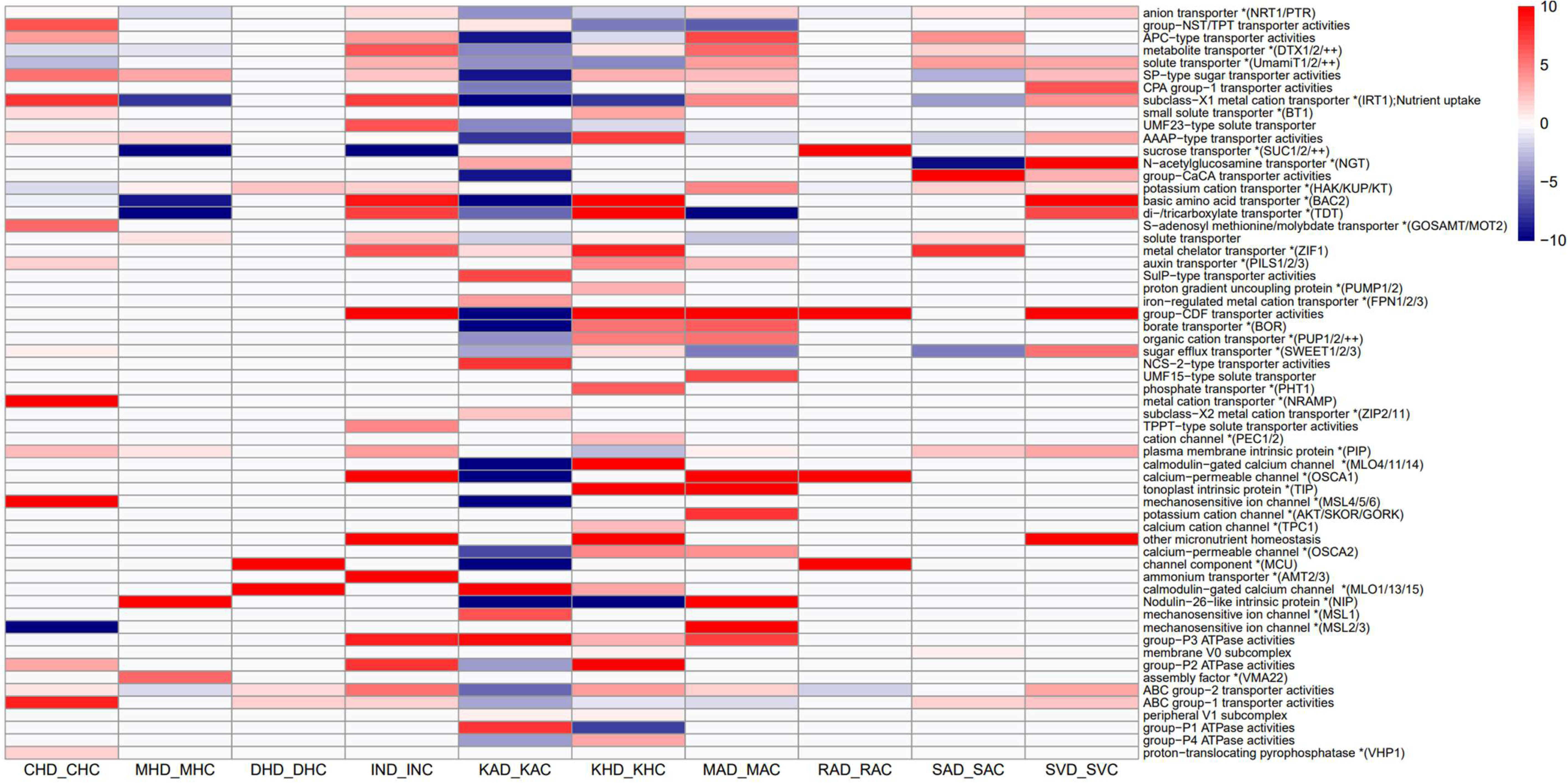
The ten durum wheat genotypes exhibit differential expression of genes involved in Solute Transport. Heatmap was built using DEGs defined as |Log2FC|>1; FDR<0.05 under Drought Stress in genotype-specific expression Patterns. The DEGs enrichment was extracted using the bin classification from Mercator4 v8.

**Fig. S11:**
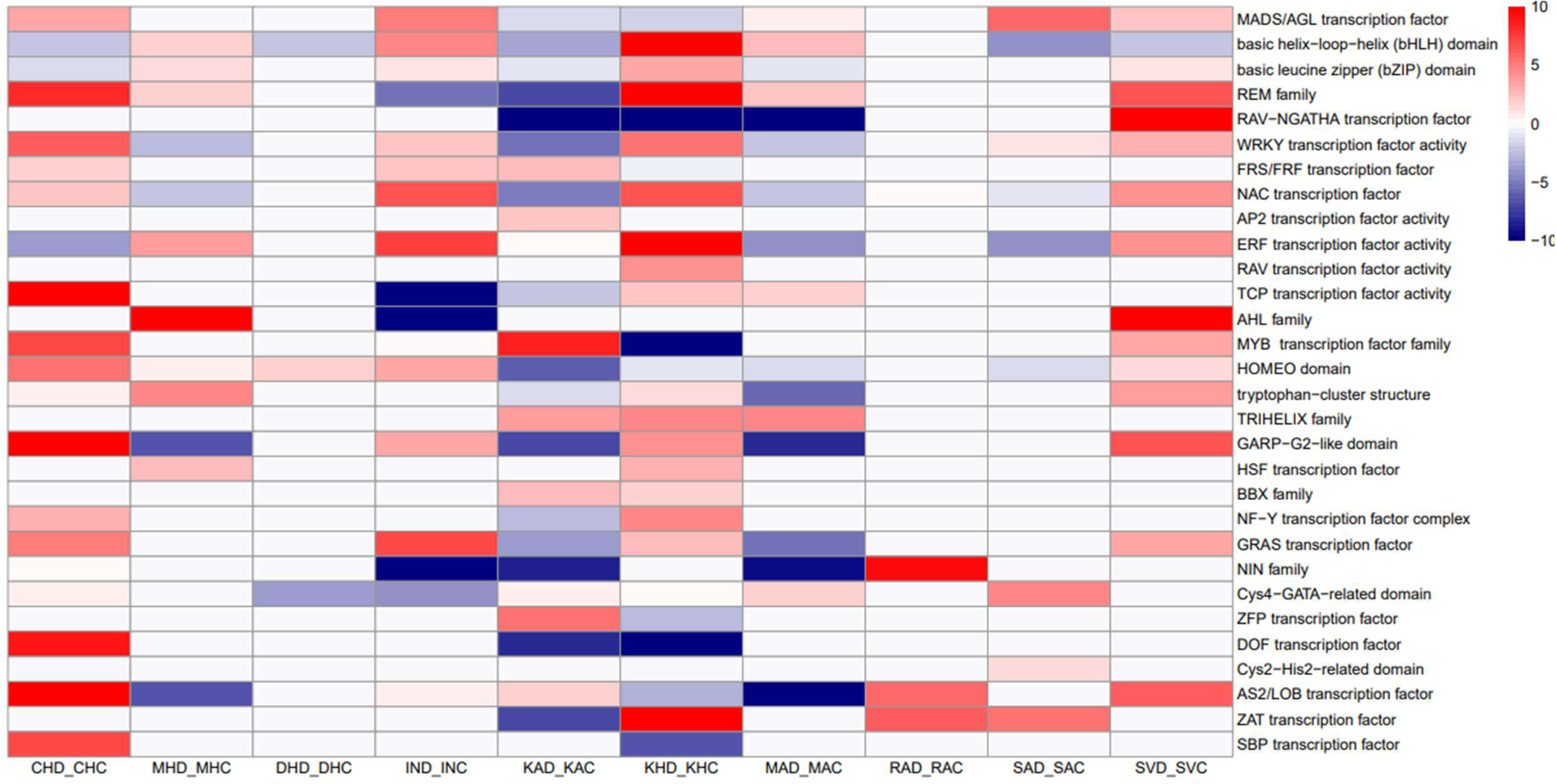
The ten durum wheat genotypes exhibit differential expression of Transcription factors. Heatmap of Transcription factors Differentially Expressed under Drought Stress in genotype-specific expression Patterns. DEGs are defined as |Log2FC|>1; FDR<0.05. The TFs <u>classes were extracted using the bin classification from Mercator4 v8.</u>

**Table S1:** List of DEGs considering |Log2FC|>1; FDR<0,05

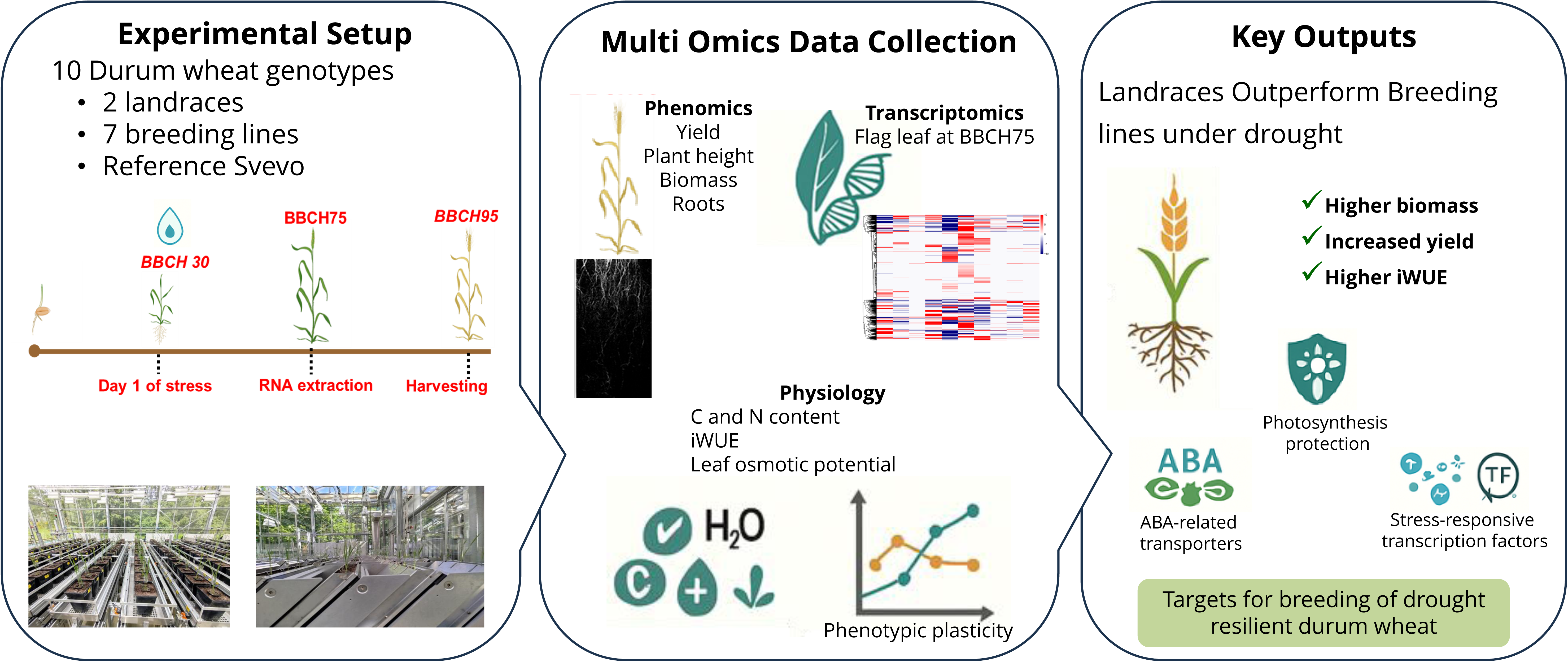

